# Rgs1 is a regulator of effector gene expression during plant infection by the rice blast fungus *Magnaporthe oryzae*

**DOI:** 10.1101/2022.09.04.506535

**Authors:** Bozeng Tang, Xia Yan, Lauren S. Ryder, Neftaly Cruz-Mireles, Darren M. Soanes, Camilla Molinari, Andrew J. Foster, Nicholas J. Talbot

## Abstract

To cause rice blast disease the filamentous fungus *Magnaporthe oryzae* secretes a battery of effector proteins into host plant tissue to facilitate infection. Effector-encoding genes are expressed only during plant infection and show very low expression during other developmental stages. How effector gene expression is regulated in such a precise manner during invasive growth by *M. oryzae* is not known. Here, we report a forward-genetic screen to identify regulators of effector gene expression, based on selection of mutants that show constitutive effector gene expression. Using this simple screen, we identify Rgs1, a regulator of G-protein signalling (RGS) protein that is necessary for appressorium development, as a novel transcriptional regulator of effector gene expression, which acts prior to plant infection. We show that an N-terminal domain of Rgs1, possessing transactivation activity, is required for effector gene regulation and acts in an RGS-independent manner. Rgs1 controls expression of at least 60 temporally co-regulated effector genes, preventing their transcription during the pre-penetration stage of development prior to plant infection. A regulator of appressorium morphogenesis is therefore also required for orchestration of pathogen gene expression required for invasive growth by *M. oryzae* during plant infection.

## Introduction

Plant pathogens secrete effector proteins into host tissues in order to suppress host immunity, modulate plant cell organisation and perturb cellular functions (1, 2). In this way, they re-programme host cells to facilitate pathogen invasion and proliferation. Effector gene expression is tightly regulated so that different families of effectors are deployed at each stage of plant infection. How plant pathogenic fungi regulate effector gene expression is, however, poorly understood. This is exemplified by the devastating rice blast pathogen *Magnaporthe oryzae* which possesses a large repertoire of effector genes expressed specifically during plant cell invasion (3). Although the function of effectors is an area of intense study (4), relatively little is known about the transcriptional control that governs effector gene expression (5–8).

In this study, we set out to investigate transcriptional regulation of effector-encoding genes in *M. oryzae* and how gene expression is orchestrated during plant infection. On the leaf surface, *M. oryzae* conidia germinate and sense plant surface cues which trigger development of a specialised infection cell, the appressorium, required for penetration of host cells (9–11). Appressorium development requires heterotrimeric G-protein signalling, to transmit surface-sensing cues to downstream modules that facilitate morphogenesis, controlled by the mitogen-activated protein kinase Pmk1 (12, 13), and cAMP-dependent protein kinase A pathways (14). G-protein subunits in *M. oryzae* are controlled by Regulator of G-protein Signalling (RGS) proteins, which are important for appressorium development (15–17).

After penetration into rice tissue, the blast fungus rapidly switches to biotrophic growth and overcomes plant immunity by secreting effector proteins (18). At the tip of the primary invasive hypha, a plant-derived membrane-rich structure, the Biotrophic Interfacial Complex (BIC), develops and remains intact as further secondary invasive hyphae fill the rice cell (19). Effectors of *M. oryzae* destined for delivery into plant cells, including avirulence gene products such as Avr-Pita, Avr-Pizt, Avr-Pii and Pwl2, all localize to the BIC (20, 21), from which they appear to be translocated and delivered into host cells (5). A second group of effectors, including Slp1 and Bas4, are secreted to the apoplast where they suppress extracellular defence responses, such as chitin-triggered immunity (22). As a consequence of effector-mediated suppression of immunity, *M. oryzae* is able to proliferate rapidly in plant tissue and move from cell-to-cell using pit fields containing plasmodesmata. The fungus develops a specialised transpressorium at rice cell junctions, which facilitates cell invasion at pit fields in a process regulated by the Pmk1 MAPK pathway (23).

A recent transcriptional profiling study has shown that effector gene expression in *M. oryzae* occurs only during plant infection. Effector genes are temporally regulated during infection with early-acting effectors expressed as soon as 8 hours after conidial germination, and large families of structurally conserved effectors, such as the Max effectors (24), expressed during biotrophic growth of the pathogen, 24-48 hours after initial infection (25). The specific temporal and spatial expression patterns suggest that *M. oryzae* effectors must be under very precise transcriptional regulation, but little is known regarding the transcriptional regulators necessary to achieve this complexity of control. In other pathogenic fungi, our understanding of effector gene regulation is also limited. In the corn smut fungus *Ustilago maydis*, for instance, a transcriptional regulator Ros1 has been implicated in spore formation and effector gene expression during the late stages of infection (26), while in the necrotrophic wheat pathogen *Parastagonospora nodorum* the Zn_2_Cys_6_ transcription factor PnPf2 positively regulates 12 effector-like protein-encoding genes (27). Recent studies have also implicated global histone modification dynamics in control of pathogen gene expression in *M. oryzae* during infection (28). There are, however, only limited reports to date that have investigated the mechanism of effector gene regulation in plant pathogenic fungi.

In this study, we set out to identify putative regulators of effector gene expression. We reasoned that because effector genes are only expressed during growth in plant tissue, it would be possible to select for mutants that exhibit constitutive effector gene expression. These would potentially carry mutations in genes encoding transcriptional regulators. Here, we report a simple forward genetic screen using a strain of *M. oryzae* in which we expressed a translational fusion of an effector, Mep2, with a green fluorescent protein tag. Using this reporter line, we selected *M. oryzae* mutants in which we could observe constitutive Mep2-GFP fluorescence in hyphae and spores. This led to identification of a mutant, *cer7*, which carries a single point mutation in a gene called *RGS1*. We show that the Rgs1 protein–which has been previously implicated as a regulator of G-protein signalling during appressorium development by *M. oryzae* (15–17) –also acts as a transcriptional regulator of effector gene expression. We provide evidence that Rgs1 is necessary for repressing the expression of at least 60 temporally co-regulated effector-encoding genes during the pre-penetration stages of development and that these genes are subsequently de-repressed during invasive growth by the fungus enabling their specific deployment in plant tissue.

## Results

### A forward genetic screen to identify regulators of effector gene expression in *M. oryzae*

In this study, we set out to identify transcriptional regulators of effector gene expression in *M. oryzae*. We reasoned that because effectors are only expressed during plant infection, selecting mutants that show constitutive expression of an effector gene, would provide a simple method to identify corresponding regulatory gene, carrying either a mutation leading to constitutive activation of a transcriptional activator, or a loss of function mutation in a repressor, for example. We therefore generated a strain of *M. oryzae* expressing an effector-encoding gene *MEP2*, fused to the Green Fluorescent Protein gene (GFP). *MEP2* was identified in a recent study that characterised the effector repertoire of *M. oryzae* based on their differential expression during pathogenesis (25). *MEP2* shows peak expression 48h after infection based on RNA-seq analysis, corresponding to a time of rapid plant tissue biotrophic colonisation by the fungus (25). We generated a *MEP2:GFP* gene fusion, transformed it into a *M. oryzae* wild type rice pathogenic strain Guy11 and selected a transformant with a single integration of the reporter gene construct. We observed specific expression and localisation of Mep2-GFP fluorescence in the Biotrophic Interfacial Complex (BIC) of invasive hyphae during rice infection, with very little detectable expression in either conidia or vegetative hyphae of the fungus (Fig. 1*A*, Fig S1). Consistent with this, quantitative real-time PCR (qPCR) showed high transcript abundance of *MEP2* in invasive hyphae (IH) with peak expression at 48 h post-inoculation (hpi) compared to minimal basal expression in conidia (Fig. 1*B*). Having established that Mep2-GFP is specifically expressed during *in planta* growth by *M. oryzae*, we carried out UV mutagenesis on conidia of the Mep2-GFP strain. We selected mutants that showed constitutive Mep2-GFP fluorescence in conidia and named them Cer (<u>C</u>onstitutive <u>E</u>ffector <u>R</u>egulator) mutants. One of these mutants, *cer7,* which has green fluorescent conidia, was selected (Fig. 1*C*). We verified constitutive expression of *MEP2* using qPCR which showed elevated expression in *cer7*, compared to the original Mep2-GFP transformant (Fig. 1*D*). We also observed constitutive expression of Mep2-GFP in mycelium grown in axenic culture, in appressoria, and in the BIC of invasive hyphae in the *cer7* mutant (*SI Appendix*, Fig. S1). Taken together, these results suggest that the expression of the *MEP2* effector gene is induced, or de-repressed in spores and mycelium of the *cer7* mutant.

**Figure 1.**
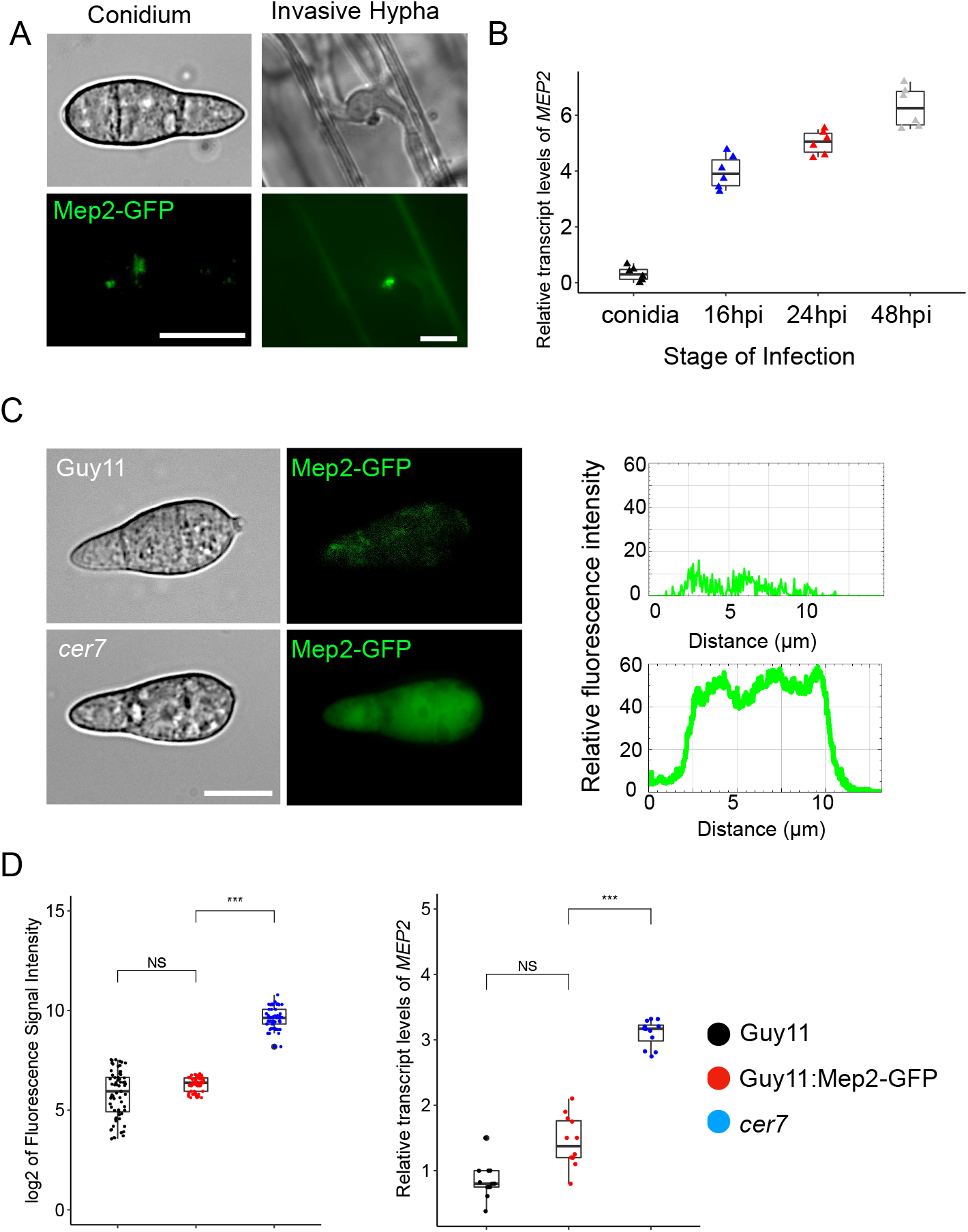
A forward genetic screen identified the *cer7* mutant of *M. oryzae* (***A***) Micrographs showing differential expression of Mep2-GFP at the BIC in invasive hyphae and basal expression in conidia of the wild type *M. oryzae strain* Guy11. (Scale bar, 10 μm). BIC localization was imaged in rice leaf sheath tissue inoculated with conidia of Guy11:Mep2-GFP at 32 hpi. (***B***) Boxplots to show relative transcripts of *MEP2* as log2 fold changes values in invasive hyphae of Guy11. The samples were harvested from Guy11 conidia and leaf sheath harvested at 16hpi, 24hpi, and 48hpi. n=6 experiments. Expression is shown relative to the *M. oryzae* actin gene. (***C***) Micrographs and line-scan graphs showing the constitutive fluorescence signal of Mep2-GFP in mutant strain *cer7* conidia, compared to the wild-type Guy11 (Scale bar, 10 μm). (***D***) Box plots to show fluorescence intensities of Mep2-GFP as log2 values and relative abundance of *MEP2* transcripts as log2 fold change values in qRT-PCR. Colours correspond to Guy11 (black), *cer7* (red), and Guy11:*MEP2-GFP* (blue). Significance between groups of samples was performed using Unpaired Student’s t-test. ***p<0.001, *p<0.05, NS = no significant difference.

### Identification of the *CER7* locus by bulked segregant analysis

To identify the mutation leading to constitutive *MEP2* expression in the *cer7* mutant we first sequenced the genome of *cer7* mutant and aligned it against the genome sequence of the original Guy11 Mep2-GFP transformant and the *M. oryzae* reference genome of strain 70-15 (29). A total of 1955 variants (SNPs and indels) were identified compared to the *M. oryzae* 70-15 reference genome sequence of which 1036 were located within the coding regions of 408 different genes. To identify the *cer7* mutation, we carried out bulked segregant analysis (30) by crossing the *cer7* (*Mat1*-*1*) mutant with a wild-type strain TH3 of opposite mating type (*Mat1*-*2*) (*SI Appendix*, Fig. S2*A*). We selected perithecia and dissected asci (*SI Appendix*, Fig. S2*B*). Ascospore progeny were then collected and phenotypically characterized based on the fluorescence signal of Mep2-GFP (Fig. 2*A*). A total of 253 progeny were selected, of which 59 progeny (23.3%) showed the *cer7* phenotype, and 194 progeny the wild type phenotype (Fig. 2*A*). The *MEP2-GFP* construct is present in a single copy in the *cer7* mutant, as confirmed by *de novo* assembly of the *cer7* genome sequence and would therefore be predicted to segregate in a 1:1 ratio in ascospore progeny. We reasoned that if *cer7* is caused by mutation at a single locus then this should also segregate in a 1:1 ratio. We would therefore expect to see the observable *cer7* fluorescent conidia phenotype in a 1:3 ratio, with a quarter of progeny showing constitutive Mep2-GFP expression, which was validated by a Chi-squared test (*χ*^2^ = 0.135, df=1, *P* = 0.713). To carry out bulked segregant analysis we then extracted genomic DNA from *cer7* and wild-type progeny, respectively, and bulked them into two separate pools for genome sequencing. This enabled us to define a region of 692kb on supercontig 8.2 which showed the highest frequency of SNPs identified in progeny showing the *cer7* phenotype (Fig. 2*B*). Within this region, only one polymorphism matched SNPs identified in the genome sequence of the *cer7* mutant, located at position 3779156 in the coding region of gene MGG_14517. This gene has previously been identified as *RGS1*, which encodes a regulator of G-protein signalling in *M. oryzae* (15). The SNP results in a single amino acid sequence change in the predicted gene product from glutamic acid to a stop codon (GAA to TAA) (*SI Appendix*, Fig. S2*C*). To confirm the association, we sequenced PCR-amplified fragments spanning the SNP in 10 *cer7* and 10 wild type progeny (*SI Appendix*, Fig. S2*D*), which verified the analysis.

**Figure 2.**
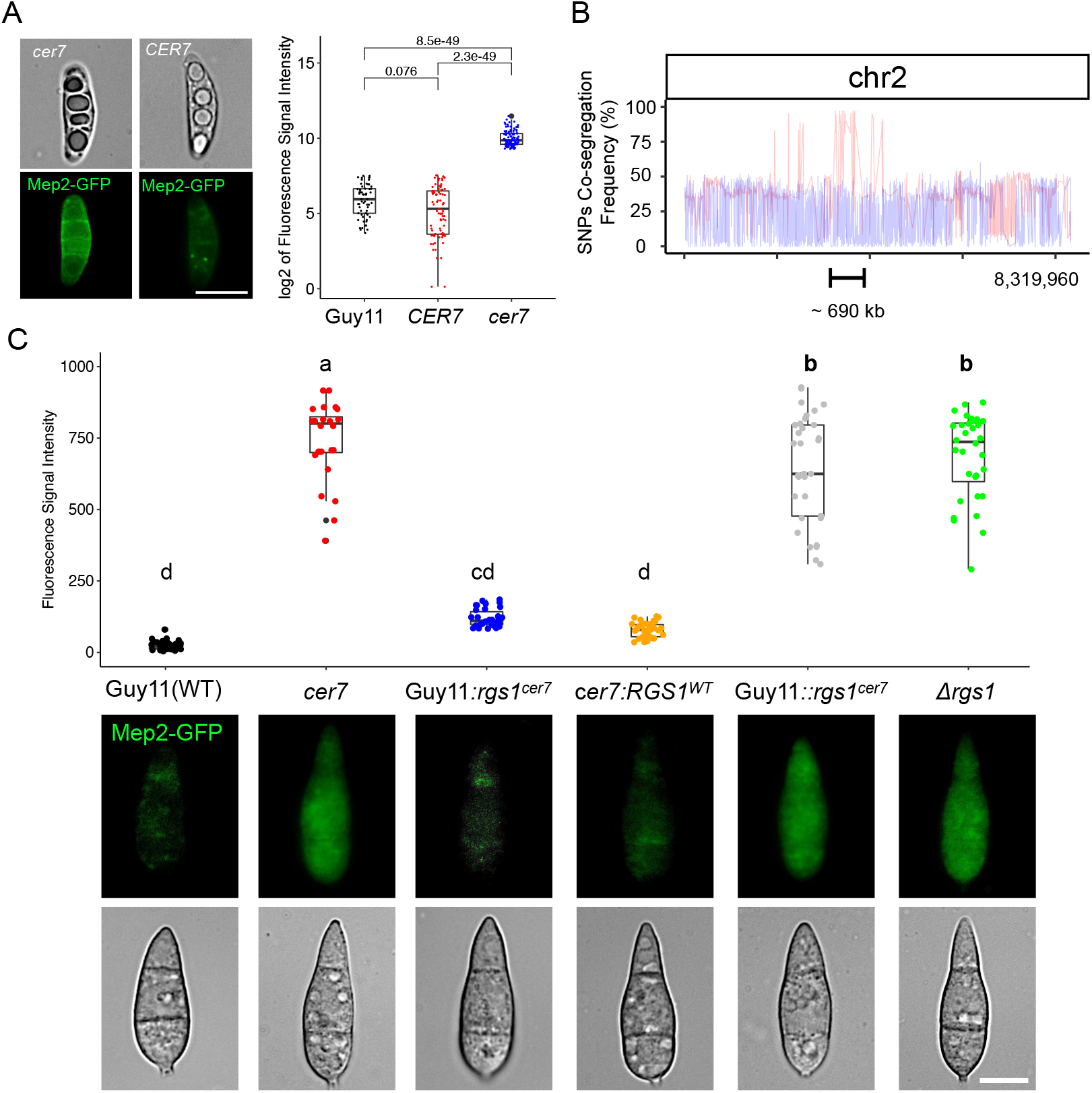
Bulked segregant analysis and genetic complementation defined *rgs1*^*cer7*^ as the allele responsible for constitutive expression of Mep2-GFP in conidia. (***A***) Micrographs showing expression of Mep2-GFP in segregating ascospores obtained from a cross between *cer7* and TH3. A single ascospore was isolated from an ascus under the dissecting microscope. Progeny displaying a high level of fluorescence signal from Mep2-GFP (*cer7*) were pooled together for whole genome sequencing. (Scale bar, 2 μm). Boxplots showing log2 values of fluorescence intensities of Mep2-GFP measured from conidia of 253 individual *M. oryzae* progeny. Progeny with *cer7* phenotype (blue), and those showing *CER7* phenotype (red), were compared to wild-type Guy11 conidia (black). n(*cer7*) =59, n(*CER7*) =194. A t-test was performed to determine the significance between the samples and *P* values are shown. (***B***) Graph showing SNP co-segregation frequencies on chromosome 2 after sequencing genomic DNA from pools of progeny segregating for *cer7* and *CER7* phenotypes, respectively. Red line shows frequencies of variants identified from pooled genomic DNA from progeny showing the *cer7* phenotype. Blue line shows frequencies of variants identified from pooled genomic DNA from progeny with *CER7* phenotype (***C***) Boxplot and micrographs to show the fluorescence intensity of Mep2-GFP in conidia of wild-type Guy11 (Black), *cer7* (red), Guy11:*rgs1*^*cer7*^ (blue), *cer7:RGS1*^*WT*^ (orange), Guy11::*rgs1*^*cer7*^ (grey), and *Δrgs1* (green). Letters within each sample refer to One-way ANOVA tests (p<0.05, Duncan test). Micrographs show bright field and epifluorescence images of conidia. (Scale bar, 10 μm).

To test whether the *RGS1* gene corresponds to *cer7* we carried out genetic complementation and allelic replacement assays. We first introduced the wild type *RGS1*^*WT*^ allele into the *cer7* mutant, which resulted in transformants with non-fluorescent conidia (Fig. 2*C*). Introducing the *rgs1*^*cer7*^ allele ectopically into the wild-type (*RGS1^+^*) Mep2-GFP strain also resulted in non-fluorescent conidia. By contrast, when we carried out targeted allelic replacement of *RGS1* with the *rgs1*^*cer7*^ mutant allele in the Mep2-GFP strain, this resulted in transformants with fluorescent conidia. When considered together, this provides evidence that *cer7* is a recessive loss of function mutation in the *RGS1* gene, consistent with the premature stop codon generated by the mutation (Fig. 2*C*). To test this idea directly, we generated a targeted gene deletion mutant in the wild type Mep2-GFP background. This led to constitutive expression of Mep2-GFP in conidia of the resulting Δ*rgs1* mutants (Fig. 2*C*). The results were validated by qPCR which confirmed that *cer7* and Δ*rgs1* mutants show high level expression of *MEP2* in conidia (*SI Appendix*, Fig. S3*A*). We also observed the Δ*rgs1* and *cer7* strains produced similar phenotypes in axenic culture, showing white aerial hyphal growth, water soaking, and aberrant appressorium formation on hydrophilic surfaces, which is consistent with previous reports of Δ*rgs1* mutants (15, 16, 31) (*SI Appendix*, Fig. S3*B*).

### Rgs1 acts as a transcriptional regulator of the *MEP2* effector gene

Rgs1 has been studied previously in *M. oryzae* as a regulator of G-protein signalling which affects asexual development, appressorium formation, surface sensing, and virulence through its interaction with the three G*α* subunit proteins MagA, MagB and MagC (15). Mutants lacking Rgs1 form appressoria on non-inductive hydrophilic surfaces and show reduced virulence resulting from mis-regulation of MagA, as well as a hyper-sporulation phenotype associated with mis-regulation of MagB (31). The reported roles of Rgs1 are therefore associated with the pre-penetration phase of development, prior to plant tissue invasion (16, 17, 31). To investigate the potential role of Rgs1 as an effector regulator, we therefore investigated the temporal expression of *RGS1* in publicly available RNA-seq data-set (PRJEB45007) (25). This showed that *RGS1* is expressed in conidia and during initial stages of appressorium formation, but then very reduced in expression during plant infection. This is the reciprocal pattern to *MEP2*, which is not expressed in conidia, but highly expressed during invasive growth, peaking in expression at 48h after infection (Fig. 3*A*). To experimentally verify this pattern of expression, we generated a *M. oryzae* transformant expressing Rgs1-GFP and carried out live cell imaging. Rgs1-GFP is highly expressed in conidia, germ tubes, and incipient appressoria, but significantly reduced in invasive hyphae (Fig. 3*B*). We then extracted total protein from mycelium and plant tissue infected with the Rgs1-GFP strain, and a control strain of *M. oryzae* expressing GFP under control of a high-level constitutive promoter ToxA (32), and performed western blot analysis with anti-GFP antibodies. We detected the predicted 106 kDa Rgs1-GFP fusion protein in mycelium, but this was not observed in infected plant tissue samples at 32h after infection (*SI Appendix*, Fig. S4*A*). To investigate the nature of Rgs1 expression further, we next analysed published RNA-seq data (PRJEB36580)(33), and found high coverage reads spanning all three exons of *RGS1* from conidia, while aligned reads from the first exon of *RGS1* were very reduced in RNA-seq data from infected plant material at 24 hpi (*SI Appendix*, Fig. S4*B*). This suggests that exon skipping of *RGS1* may occur during plant infection, resulting in a lower abundance of the N-terminal domain of the Rgs1 protein. It has previously been reported that Rgs1 undergoes endoproteolytic cleavage which leads to generation of a tandem Dishevelled, Egl-10, Pleckstrin domain (DEP-DEP) protein from the N-terminus of Rgs1 (N-Rgs1), encoded by the first and second exon, and a separate RGS core domain protein from the C-terminus (C-Rgs1) (31). It has been proposed that the N-Rgs1 protein is required for vesicular membrane targeting of the protein, while the C-Rgs1 protein is sequestered in the vacuole, providing a post-translational mechanism to regulate the catalytic activity of Rgs1 on its G*α* sub-unit substrates. Given the low level of N-Rgs1 associated transcripts during plant infection, we wondered whether the effector regulation function resided in N-Rgs1 and occurred during the pre-penetration phases of development. To investigate whether N-Rgs1 can act as a transcription factor, we therefore tested its transactivation activity and DNA-binding ability in yeast. We found that when N-Rgs1 is fused to the Gal4 DNA-binding domain it is able to act as a transcriptional activator (Fig. 3*C*), but when fused to the Gal4 activation domain, it is unable to bind DNA (*SI Appendix*, Fig. S5*A*). Meanwhile, neither the full length Rgs1 nor the C-Rgs1 protein show any transactivation or DNA-binding activity (*SI Appendix*, Fig. S5*A*). These results suggest that N-Rgs1 might function independently to regulate *MEP2* transcription, consistent with the position of the *cer7* premature stop codon mutation and the potential that exon skipping of Rgs1 takes place during plant infection. To test this idea, we constructed vectors carrying sequences encoding N-Rgs1 or C-Rgs1 respectively, driven by the native *RGS1* promoter and terminator sequences, and transformed these into the *M. oryzae cer7* mutant. We found that N-Rgs1 was able to complement *cer7* preventing expression of Mep2-GFP in conidia (Fig. 3*D*). By contrast, expressing C-Rgs1 did not complement the *cer7* phenotype and conidia remained fluorescent (Fig. 3*D*). In control experiments we verified *cer7* complementation with the full-length *RGS1* gene and lack of complementation with the *RGS1^cer7^* allele. When considered together, these results suggest that N-Rgs1 acts as a repressor of transcription of *MEP2* in conidia, preventing its expression prior to plant infection. However, given that N-Rgs1 is unable to bind DNA, which was also confirmed using a yeast-one hybrid assay which did not find any evidence for N-Rgs1 binding to the *MEP2* promoter (*SI Appendix*, Fig. S5*B*), it is likely that N-Rgs1 regulates transcription of *MEP2* indirectly, perhaps by activating a repressor protein, or acting in association with another partner to bring about *MEP2* repression. We conclude that N-Rgs1 is necessary for regulation of the *MEP2* effector gene.

**Figure 3.**
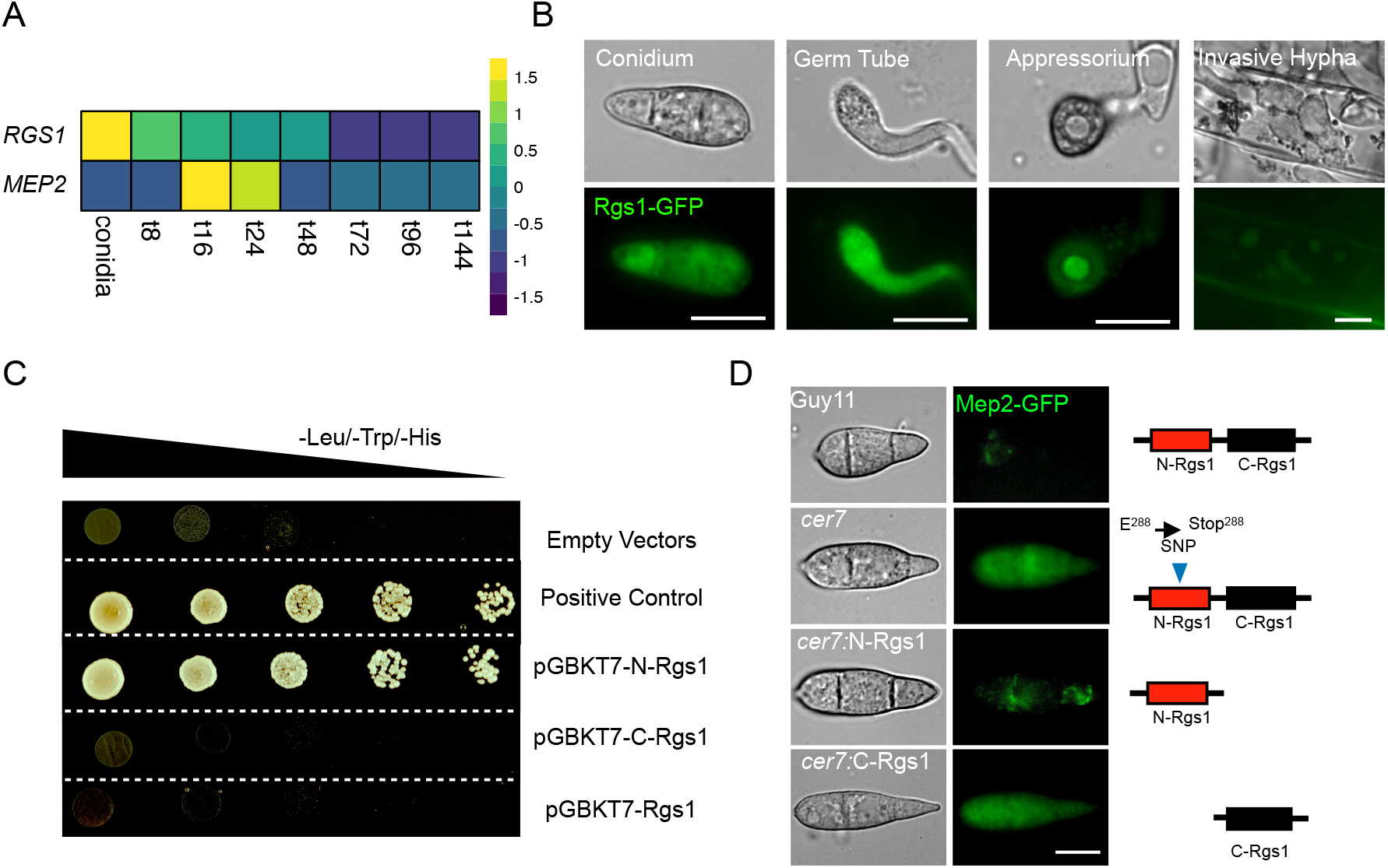
The N-terminus of N-Rgs1 is required for the repression of *MEP2* expression in conidia. ***A***) Heatmap showing relative transcript abundance of *MEP2* (MGG_00230) and *RGS1* (MGG_14517) genes in *M. oryzae* conidia, and during infection from 8 to 144 hpi. Relative transcript levels are fold change compared to expression in mycelium (from data set PRJEB44745). Data were extracted from RNA-seq dataset PRJEB45007. The colour key shows scaled fold change values. (***B***) Micrograph showing the fluorescence signal of Rgs1-GFP expressed in Guy11. Live-cell imaging was performed during a time course experiment to investigate expression of Rgs1-GFP in conidia, germ tubes, mature appressoria, and invasive hyphae at 24hpi. (***C***) Images showing transactivation activity of N-terminal Rgs1 (N-Rgs1), C-terminal Rgs1 (C-Rgs1), and full length Rgs1 in yeast cells. Co-transformation of Y2HGold yeast cells with bait (BD) and prey (AD) vectors was carried out with following combinations; pGBKT7-N-Rgs1 / pGADT7, pGBKT7-C-Rgs1 / pGADT7, pGBKT7-Rgs1/ pGADT7, and positive control (pGBKT7-53 and pGADT7-T) along with empty vectors, and grown in double drop-out and quadruple-dropout media. Images represent two independent biological replicates. (***D***) Micrographs showing expression of Mep2-GFP in conidia of strains *cer7*, Guy11, *cer7*:N-Rgs1, *cer7*:C-Rgs1. Conidia from each strain were harvested from colonies after 5 days growth on CM and immediately mounted on microscope slides for GFP visualization using epifluorescence microscopy. (Scale bar, 10 μm). The schematic illustration demonstrates different genotypes in corresponding *M. oryzae* strains.

### Rgs1 regulates effector gene expression independently of its G-protein signalling function

Rgs1 has been reported to accelerate the intrinsic GTPase activity of target Gα subunits during appressorium development (15) and we therefore decided to test directly whether its RGS activity, which resides in the C-Rgs1 part of the protein, was associated with its ability to regulate *MEP2* transcription. We decided to test whether repression of *MEP2* transcription during the pre-penetration stage of development is regulated via the action of Rgs1 on its associated Gα-subunits in response to surface cues, thereby enabling Gα subunits to activate their downstream effector modules to trigger *MEP2* transcription. We reasoned that if *MEP2* transcription is G-protein-dependent, then mutants affected in their sensitivity to Rgs1 signalling, lacking GTPase activity, or constitutively activated GTP-Gα-subunits, would be affected in *MEP2* transcription. To test this idea, we generated Rgs1-insensitive alleles MagA(G187S), MagB(G183S), and MagC(G184S), the GTP-ase inactive MagA(Q208L) and MagB(Q204L) alleles, and the MagB(G42R) constitutively activated allele (15), and introduced them each into the Guy11 strain expressing Mep2-GFP. We did not observe increased Mep2-GFP expression in conidia of any of these transformants, suggesting that Rgs1 regulates transcription of *MEP2* in a G-protein-independent manner (*SI Appendix*, Fig. S5*C*). Taken together, these results provide evidence that the N-Rgs1 protein is necessary to regulate transcription of *MEP2* in conidia and acts independently of the G-protein signalling role of Rgs1.

### Rgs1 regulates a large group of effector genes in *M. oryzae*

To determine the wider function of Rgs1 we tested the ability of the Δ*rgs1* and *cer7* mutants to cause rice blast disease. We inoculated 21-day-old seedlings of the susceptible rice cultivar CO-39 and scored disease symptoms after 5 days. A significant reduction in disease lesion number was observed in Δ*rgs1* and *cer7* infections compared to the isogenic wild type Guy11 (Fig. 4*A*). Rgs1 has previously been reported to play a role in pathogenesis, but this has been attributed to its G-protein regulatory function (11). We decided to test whether Rgs1 regulates a wider group of virulence-associated genes by performing comparative global transcriptional profiling using Δ*rgs1* and *cer7* mutants compared to Guy11. We reasoned that genes regulated by Rgs1 during plant infection would show a similar de-repression in conidia of Δ*rgs1* and *cer7* mutants. We therefore extracted mRNA from conidia Δ*rgs1*, *cer7*, and Guy11 using a total of five biological replicates of the experiment and performed RNA-seq analysis. Euclidean analysis was used to determine the similarity between expression profiles in each mutant, revealing a strong overlap between Δ*rgs1* and *cer7* mutants (Fig. 4*B*). This was consistent with principal component analysis which also demonstrated very similar transcriptional patterns between the *cer7* and Δ*rgs1* mutant (*SI Appendix*, Fig. S6*A*). In total, 996 genes were upregulated in conidia of *cer7*, and 1126 in Δ*rgs1* compared to Guy11 (log2|FC|>1, padj<0.05). Of these, 757 are shared between *cer7* and Δ*rgs1*. Metabolic enrichment analysis of the Rgs1-repressed gene set showed over-representation of gene functions associated with starch and sucrose metabolism, and glycan degradation, which are also induced during biotrophic invasive fungal growth, as well as phenylalanine and tyrosine metabolism which may be associated with secondary metabolism (*SI Appendix*, Fig. S6*B*). We also observed over-representation of biological processes associated with membrane function and transmembrane transport, as well as oxido-reductases. These functions too are associated with biotrophic invasive growth of *M. oryzae* (25) (*SI Appendix*, Fig. S6*C*). A recent study of the transcriptional landscape of plant infection by *M. oryzae* used weighted gene co-expression network analysis (WCGNA) to define temporal co-expression clusters of *M. oryzae* genes during infection (25) (*SI Appendix*, Fig. S7*A*). We observed that Rgs1-regulated genes can be classified into each WCGNA cluster but genes upregulated in conidia of Δ*rgs1* mutants are enriched in cluster M5, which peaks in expression at 48 h after infection (*SI Appendix*, Fig. S7*A*). This cluster also contains many effector-encoding genes (25). Consistent with this, we found 98 predicted or known effector-encoding genes upregulated in conidia of Δ*rgs1* mutants, of which 60 were also up-regulated in conidia of the *cer7* mutant (Fig. 4*C*) (*SI Appendix*, table S3). This suggests that in addition to *MEP2*, Rgs1 may regulate a much larger group of effectors, that are de-repressed in conidia when the function of Rgs1 is compromised. Among this group, there were 8 previously reported effector candidates, including *BAS3*, *BAS113*, *MEP19*, and *MEP27* (8, 25), and the necrosis and ethylene-inducing (Nep-like) peptide effector *NLP4* which is involved in programmed cell death in plant tissue (34) (Fig. 4*D*). To test whether the identified effectors are also de-repressed in a *cer7* mutant we generated *BAS113-RFP* and *BAS3-RFP* gene fusions and transformed them into the *cer7* mutant strain and Guy11, respectively. We observed high levels of Bas3-RFP and Bas113-RFP fluorescence in conidia of the *cer7* mutant transformants, compared to Guy11 (Fig. 4*E*), which provides further evidence that Rgs1 regulates additional effectors to Mep2. Transcriptional profile analysis of the 60 putatively Rgs1-repressed effectors during infection demonstrated that they peak in expression 48 and 72hpi (*SI Appendix*, Fig. S7*B*).

**Figure 4.**
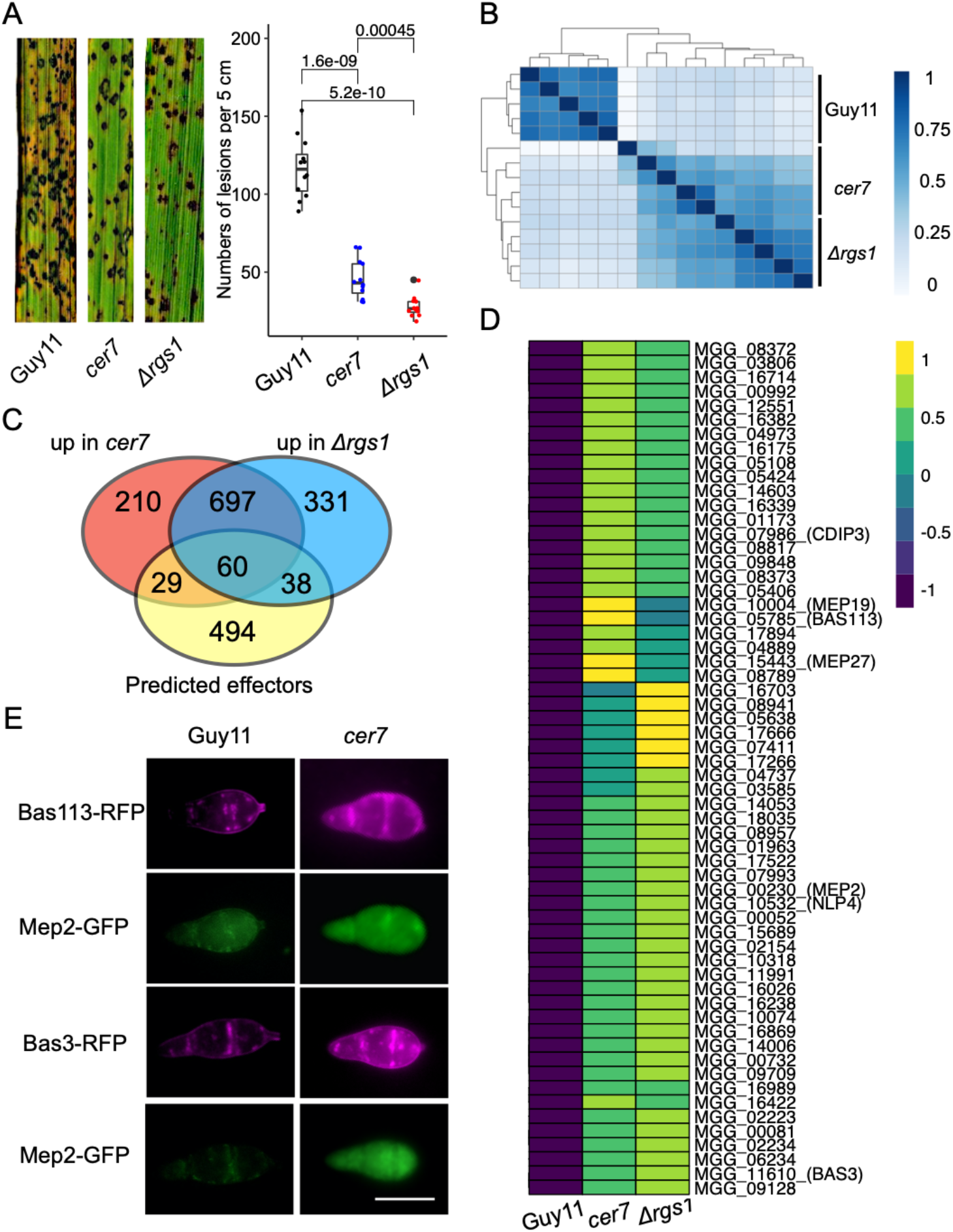
Rgs1 Regulates expression of a sub-population of effectors during plant infection. (***A***) Seedlings of rice cultivar CO-39 were inoculated with *M. oryzae* conidial suspensions of equal concentration (1 × 10^5^ conidia/mL) of wild-type Guy11, *cer7* and *Δrgs1* mutants. The boxplot represents the number of rice blast disease lesions per 5 cm in three independent repetitions of the experiment. Unpaired Student’s t-test was performed to determine significant differences. (***B***) Heatmap showing the Euclidean distance between RNA-seq samples from conidia of *cer7*, the wild type Guy11, and *Δrgs1*. Normalized reads counts were used from all the samples to determine clustering. Intensity of colours represent similarities and distance between samples. (***C)*** Venn diagram to show the number of effector genes de-repressed in conidia of the *cer7* and *Δrgs1* mutants. Blue circle = *cer7*, red circle =*Δrgs1*. (**D**) Heatmap showing expression of 60 effector genes significantly upregulated in conidia of *cer7* and *Δrgs1* mutants, compared to Guy11. Normalized expression values of transcripts used the TMM method. (***E***) Micrographs showing expression of Mep2-GFP, Bas113-RFP, and Bas3-RFP in conidia of *cer7* compared to Guy11. Conidial suspensions from each strain were inoculated onto hydrophobic glass coverslips and imaged using epifluorescence microscopy. (Scale bar, 10 μm).

### Rgs1 contributes to pathogen fitness during rice blast disease

Given that Rgs1 potentially regulates the expression of a large group of effectors, we decided to investigate the biological significance of the effector regulation mediated by Rgs1. When we inoculated rice leaf sheath with conidia of Δ*rgs1*, we found that invasive hyphae grew more slowly than those of Guy11 (Fig. 5*A*). We therefore tested whether this reduced growth in plant tissue was a consequence of the mis-regulation of effector gene expression. As many effectors act to suppress plant defence, we performed a qPCR experiment to investigate expression of a sub-set of defence-related rice genes from leaf sheath samples following infection by Guy11 and Δ*rgs1*, respectively. Transcripts of rice *PR1a* (35) and *CPS2* (36), are induced significantly at 16hpi and at 24hpi in rice tissue inoculated by strain Δ*rgs1*, compared to infection with Guy11 or non-infected tissue (Fig. 5*B*). These results are consistent with Rgs1 being required for the correct temporal dynamics of effector gene expression. However, we recognised that the role of Rgs1 was most likely to be involved in repressing effector gene expression prior to plant infection. We therefore decided to investigate the consequence of over-expressing *RGS1* throughout plant infection. For this we generated a ToxAp*:RGS1* construct and introduced it into the Mep2-GFP background. Over-expression of Rgs1 led to complete repression of Mep2-GFP during invasive growth with no visible BIC localisation observed in Guy11 (*SI Appendix*, Fig. S8). We then tested the ability of the *ToxAp:RGS1* strain to cause rice blast disease (Fig. 5*C*). We did not find a clear reduction in disease symptoms in lesion density so we decided to use a recently described relative fitness assay to evaluate the effect of over-expressing Rgs1 (25). For this, we used a mixed spore inoculum of the *ToxAp:Rgs1-GFP* strain and a wild type Guy11 strain expressing H1-RFP. The fluorescent markers are simply used as a means of visually distinguishing conidia of each strain. Conidia were mixed in a 1:1 ratio and used to inoculate CO-39 seedlings (Fig. 5*D*). We allowed disease symptoms to develop and then recovered conidia from lesions. We recorded the ratio of each spore type (green or with red nuclei, respectively) recovered from disease lesions and then carried out a second round of infection using the same ratio. We observed that the proportion of *ToxAp:RGS1* conidia reduced after two generations of infection to 30.83% and showed a fitness coefficient of 0.45 after two generations (Fig. 5*E*). We conclude that over-expression of *RGS1* has an important fitness consequence for *M. oryzae* consistent with its action as a transcriptional regulator of genes associated with biotrophic fungal growth.

**Figure 5.**
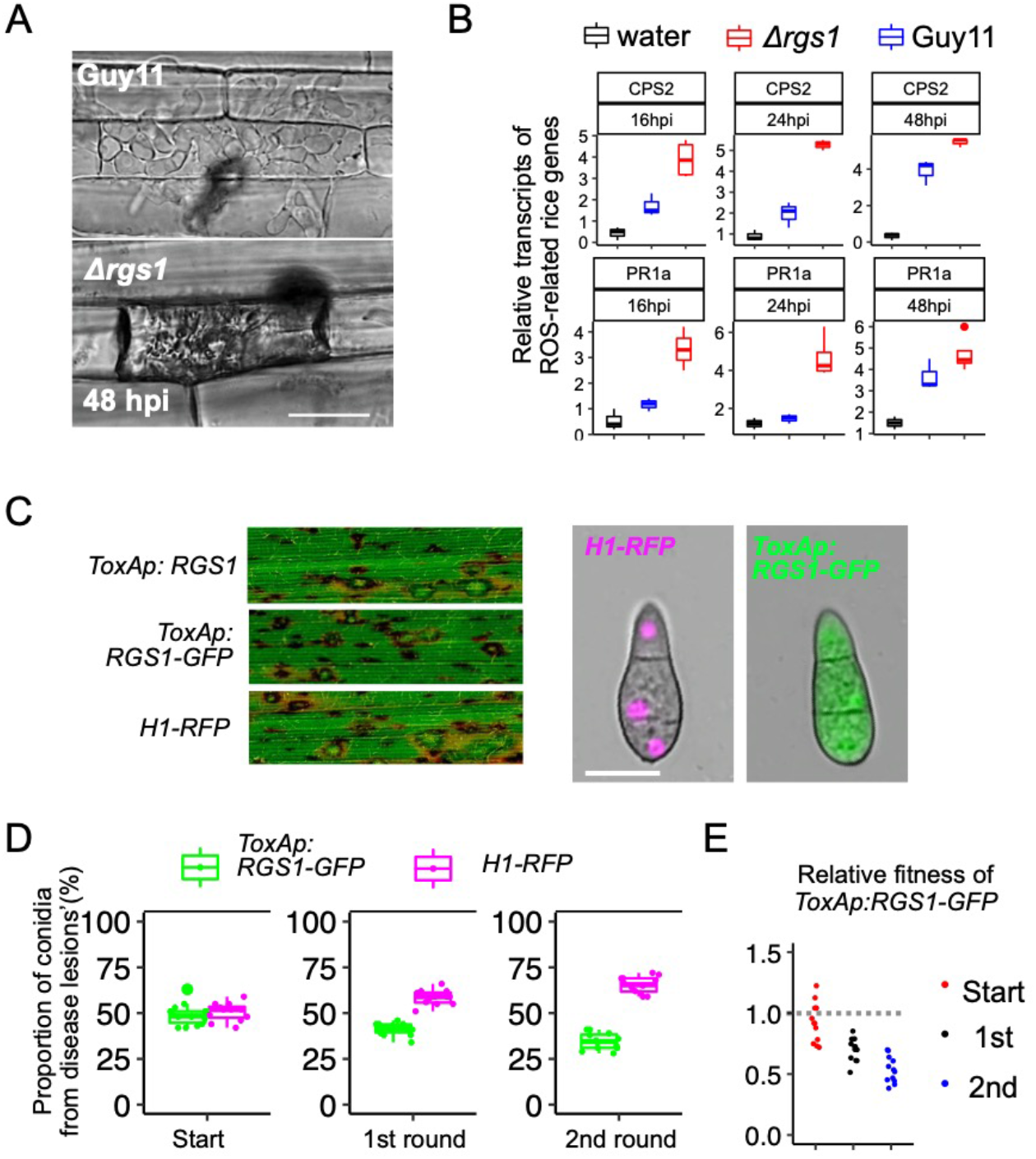
Rgs1 prevents rice defence gene expression and is a fitness determinant for rice blast disease. (***A***) Micrographs showing growth of invasive hyphae of Guy11 and *Δrgs1* 48hpi. (Scale bar, 10 μm). Rice leaf sheaths of rice cultivar CO39 were inoculated with conidial suspensions of Guy11 and *Δrgs1*. (Scale bars,12 μm). (***B***) Boxplots to show fold change values as relative transcripts of defence-related genes *CPS2* and *PR1a* in rice. Rice leaf sheath samples were inoculated with 0.02% gelatin (black), conidial suspension of Guy11 (blue) and *Δrgs1* (red). The qRT-PCR was performed using rice housekeeping genes eEF1A and UBQ5 as standards. (***C***) Seedlings of rice cultivar CO-39 were inoculated with conidial suspensions of equal concentration (1 × 10^5^ conidia/mL) of *ToxAp:RGS1*, *ToxAp:RGS1-GFP* and Guy11:H1-RFP. Micrographs show representative images of blast disease symptoms of each strain. Conidia were harvested from colonies after 5 days growth on CM and immediately mounted on microscope slides for GFP visualization using epifluorescence microscopy. (Scale bar, 10 μm). (***D***). Boxplots showing the number of spores recovered from disease lesions following mixed infections with conidial suspensions of *H1-RFP* and *ToxAp:RGS1-GFP*. After 7 days, spores were collected from disease lesions, the ratio of each genotype determined and then used for subsequent inoculation of CO-39 seedlings. (***E***). The relative fitness of *ToxAp:Rgs1-GFP* was carried out using the formula: Relative fitness = x_2_(1-x_1_) / x_1_ (1-x_2_), where x_1_ is the initial frequency and x_2_ the final frequency.

## Discussion

One of the hallmarks of fungal effector proteins is that they are expressed specifically during pathogenesis. Indeed, one of the main strategies for identifying putative effector candidates has been to use transcriptional profiling to identify differentially expressed pathogen genes. This has revealed the presence of temporally co-expressed families of effector genes in many fungal pathogens including *Ustilago maydis* (37) and *M. oryzae* (25). In rice blast, a very large repertoire of 546 effector genes has been reported, which are temporally co-expressed throughout invasive growth between 8h and 144h after infection, with large families of structurally-related effectors co-expressed between 24h and 48h after infection during biotrophic proliferation of the fungus in rice tissue (25). The complexity of the spatio-temporal regulation of effector gene expression–in which cytoplasmic effectors localise specifically at the BIC during invasive hyphal growth –suggests that sophisticated transcriptional regulation must be present to ensure that effectors are produced and deployed at the correct time and in the correct cells. But even though effectors have been studied intensively in *M. oryzae*, leading to new insight into their host targets and structure-function relationships (38, 39), very little is known about effector gene regulation. In contrast to bacterial pathogens, where effector gene regulation is well understood and associated with co-ordinate operon-based gene control (40), there have been very few regulators of fungal effector expression identified to date.

In this study, we developed a very simple forward-genetic screen to identify potential regulators of effector gene expression in *M. oryzae*. Using a reporter line of *M. oryzae*, we selected mutants showing constitutive expression of an effector-encoding gene, by simply picking mutants with green fluorescent spores–a stage of development when effectors are not normally expressed. Using whole-genome sequencing and bulked segregant analysis, we identified Rgs1, a well-known regulator of G-protein signalling during appressorium development, as being necessary for expression of the effector-encoding gene *MEP2*. Rgs1 is necessary for the repression of transcription of a group of effector-encoding genes in conidia, that are among a group of 697 genes which are Rgs1-dependent. This repression is clearly important biologically, as over-expressing Rgs1 during plant infection leads to a reduction in pathogen fitness.

Rgs1 is one of a group of 8 RGS proteins in *M. oryzae* that play distinct and overlapping regulatory functions (17). Rgs1 is involved in mediating perception of a hard, hydrophobic surface to induce appressorium morphogenesis. The non-canonical G-protein coupled receptor (GPCR), Pth11, responds to this inductive cue, leading to dissociation of the G*α*-sub-unit, MagA, from the heterotrimeric G*αβγ* complex (17). MagA activity is regulated by Rgs1, to control the levels of cAMP, leading to activation of the cAMP/protein kinase A pathway (17, 31). In addition, Rgs1 negatively regulates the G*α*-sub-unit protein, MagB, during asexual conidiation (15) and mating (16). Rgs1 is phosphorylated by casein kinase 2 at the plasma membrane and late endosome, which is essential for its GTPase-activating protein (GAP) activity (17). RGS proteins are well known to mediate GPCR signalling functions due to DEP-domain-mediated tethering (41), which is consistent with the tandem DEP domains found in Rgs1 (31). An important question arising from this work is how an RGS protein, such as Rgs1, could also exert a transcriptional regulator function. It is not clear, for instance, whether this is a direct function of Rgs1 or whether the RGS protein regulates expression of additional transcription regulators via a cell signalling function. Previously, it was reported that Rgs1 undergoes endoproteolytic cleavage to produce the DEP-DEP protein N-Rgs1, and the RGS catalytic protein C-Rgs1 (15). This mirrors the endoproteolytic cleavage of the Sst2 RGS protein in *Saccharomyces cerevisiae*, which has been proposed to serve a regulatory function of its RGS catalytic activity (41, 42). Interestingly, we found evidence in RNA-seq data from a time-course of rice infection of a potential exon skipping event that may occur in *M. oryzae*, suggesting that N-Rgs1 is preferentially generated during the pre-penetration stage of development. Furthermore, the DEP-DEP N-Rgs1 protein is able to complement the Δ*rgs1* mutant phenotype, while the RGS C-Rgs1 cannot. Additionally, Rgs1-insensitive alleles of MagA, MagB and MagC, or GTPase-inactive or constitutive alleles of MagA and MagB, showed no effect on the effector-regulatory ability of Rgs1. When considered together, this is consistent with a model whereby the effector regulatory function of Rgs1 resides in N-Rgs1 and is associated with the pre-penetration stage, when Rgs1 regulates MagA and cAMP levels during appressorium morphogenesis (15, 17, 31). Interestingly, N-Rgs1 also shows activity in a yeast transactivation assay. However, the absence of DNA-binding activity in N-Rgs1, suggests that if the protein does have a transcriptional regulatory function then it must act in association with other proteins. There are examples of RGS proteins exerting transcriptional functions (43). In humans, for example, RGS2, RGS10, and RGS12, are nuclear proteins, while RGS4, RGS14 and RGS16 are nucleocytoplasmic shuttle proteins (44, 45). Human RGS6 is, furthermore, subject to complex alternative splicing with 36 distinct splice variants present that either localise to the cytoplasm, nucleus or nucleolus of neurons in the brain (46). RGS12, meanwhile, has been shown to have transcriptional activity, which resides solely in an N-terminal domain which can act as a transcriptional repressor and also has cell cycle-regulating activities, independent of its RGS domain (43). In our experiments Rgs1 appears to localise to the cytoplasm predominantly with some punctate distribution also observed, although N-Rgs1 has also been reported to show endomembrane/vesicular localisation (31). It is therefore possible that Rgs1 interacts with the plasma membrane or endomembrane compartments via its DEP domains and exerts a signalling function that ultimately activates a downstream transcription factor (47). The RGS protein FlbA in *Aspergillus niger*, for example, involved in regulation of sporulation regulates expression of many downstream functions via a set of transcription factors, including rpnR which regulates protein secretion and stress responses (48), and Fum21 which regulates production of the mycotoxin fumonisin (49). We are currently screening putative interactors of N-Rgs1 to identify its potential mode of action in the regulation of effector function and investigating whether a sub-population of N-Rgs1 might enter the nucleus.

In summary, we have demonstrated a successful way to identify a novel regulator of effector gene expression. By using a simple forward genetic screen, it has proven possible to identify a regulator that ensures the correct temporal expression profile of a large group of at least 60 effector genes, preventing their premature expression prior to plant infection. This genetic approach has therefore revealed an unexpected link between the developmental biology of the pre-penetration stage of *M. oryzae*, regulated by the RGS protein Rgs1, and events that occur after host cell invasion, including the expression of fungal effectors that suppress plant immunity and contribute to rice blast disease.

## Materials and Methods

### Fungal and plant growth conditions

Growth and maintenance of *M. oryzae* isolates, media composition, nucleic acid extraction, and fungal transformation were all performed as previously described (50). Strains were collected from stocks and inoculated to solid complete medium (CM) and incubated at 24°C with a 12 hour light-dark cycle. For plant infection, conidial suspensions of *M. oryzae* at 5 ×10^4^ conidia mL^−1^ in 0.2% gelatin were spray inoculated onto 21-day-old rice seedlings of CO39 using an artist’s airbrush. Rice blast symptoms were scored using disease lesion density 5 days post-inoculation, as described previously (50). DNA amplification, cDNA synthesis, and qRT-PCR were carried out using standard procedures with specific primers (*SI appendix*, Table S1). Further details regarding *M. oryzae* mutant generation, UV mutagenesis, and bulked segregant analysis are provided in *SI Materials and Methods*.

### Appressorium development assays and leaf sheath infection

Appressorium assay of rice blast fungus was performed as previously described (33). Briefly, conidial suspensions were prepared at 5 ×10^4^ conidia mL^−1^ in water, and 20 μL of the suspension placed onto the surface of a hydrophobic coverslip (Corning) and incubated in a controlled environment chamber at 24°C. Live cell imaging of plant penetration, effector localization in invasive hyphae and growth of the fungus *in planta* used a leaf sheath assay, described previously. Briefly, a suspension of 5 ×10^4^ conidia mL^−1^ was prepared in 0.2% gelatin and inoculated into the hollow space of the leaf sheath tissue that was dissected from the flag leaf of 21-day-old rice seedlings of cultivar CO-39. A single epidermal layer of the leaf sheath was trimmed and mounted for microscopy (23).

### Live Cell Imaging and Quantitative Analysis

To screen the mutant, Sterile water was used to harvest spores from a 7-day old *M. oryzae* CM plate culture, before filtering through sterile Miracloth (Calbiochem). A 20 μL aliquot of suspension was transferred to a microscope slide and covered with a clean cover slip before imaging with epifluorescence and differential interference contrast (DIC) microscopy. For epifluorescence and differential interference contrast (DIC) microscopy, samples were observed using an IX-81 inverted microscope (Olympus) and a UPlanSApox100/1.40 oil objective. Images were captured using a photometrics CoolSNAP HQ2 camera system incorporated with MetaMorph software packages (MDS Analytical Technologies, Winersh, UK). Epifluorescence parameters for GFP were excitation 488 at nm, and RFP excitation at 561nm. Quantification of fluorescence intensity was performed using Fuji-ImageJ (51). To measure fluorescence intensity, conidia of interest were selected using drawing tools. “Measure” was used to obtain the “Integrated Density” (ID) for the conidial fluorescence signal. The background region next to the conidium of interest was selected and measured to obtain the pixel value for “mean fluorescence of background readings” (MFBR). The values of conidia and background were extracted and saved to calculate the corrected cell fluorescence (CCF) using the formula: CCF = Integrated Density (ID) - mean fluorescence of background readings (MFBR). Details of RNA extraction, RNA-seq analysis, quantitative real-time PCR, yeast transactivation assay, and one-yeast hybrid system are given in *SI Materials and Methods*.

## SI Materials and Methods

### Generation of *M. oryzae* mutants and strains expressing GFP or RFP fusions

To generate *M. oryzae* transformants expressing Mep2-GFP, the full-length coding sequence of *MEP2* (25) was cloned with 1.5 kb of upstream promoter sequence and fused at its C-terminus to eGFP. The Mep2-GFP plasmid was constructed using the linearized pCB1532 carrying the acetolactate synthase gene which confers resistance to sulfonylurea. Independent *M. oryzae* transformants were used for screening and selected on medium with sulfonylurea (400 μg mL^−1^) for fluorescence localization and quantification. In-Fusion cloning (Clontech Laboratories) was used to generate *RGS1^WT^*, *rgs1^cer7^*, N-Rgs1, C-Rgs1, Bas113-RFP, Rgs1-GFP, Bas3-RFP, *MagA^G187S^*, *MagB^G183S^*, *MagA^Q208L^*, *MagB^Q204L^, MagC^G184S^*, *MagB^G42R^*. *MagA^G187S^*, *MagB^G183^*^S^, and *MagC^G184S^*, and *pToxA-Rgs1-GFP* strains, using primers in Table S1. *RGS1^W^*^T^, *rgs1^cer7^*, N-Rgs1, C-Rgs1, were cloned under control of the *RGS1* native promoter (1 kb) and native terminator (600bp) sequence, except N-Rgs1 which was fused to the TrpC terminator., Bas113-RFP, Rgs1-GFP, Bas3-RFP, and pToxA-Rgs1-GFP were cloned with their full protein coding sequences without stop codons driven by their native promoters, except *pToxA-Rgs1-GFP* which was fused with ToxA promoter at its N-terminus. The fragments were then fused with eGFP in-frame at the C-terminus. *MagA^G187S^*, *MagB^G183S^*, *MagA^Q208L^*, *MagB^Q204L^, MagC^G184S^*, *MagB^G42R^*. *MagA^G187S^*, *MagB^G183^*^S^, and *MagC^G184S^* were cloned with their native promoters and point mutations generated as described previously (15). Constructs were routinely cloned into a SpeI and NotI digested StrataClone vector, pSC-A-amp/kan, with the BAR gene conferring bialophos resistance (52, 53). Plasmids were then introduced into *M. oryzae* by fungal transformation (54) and transformants selected by growing on CM supplemented with glufosinate (30 μg mL^−1^). All transformants were evaluated by PCR using primers flanking the predicted insertion to select those carrying single plasmid insertions. This was subsequently verified by whole genome sequencing.

Targeted gene replacement of the Rgs1-encoding gene and native allelic replacement of *RGS1* with the hygromycin phosphotransferase gene cassette (HYG), bestowing hygromycin B resistance, were performed using the split-marker strategy as previously described with minor modifications (55–57). Briefly, we first amplified the 1 kb *HY* and 0.7 kb length YG fragments using the primers M13F and HY split for *HY*, M13R and YG split for *YG* (55). To create the fusion in the first round of PCR, a 3.2 kb length fragment of DNA sequence of the mutant allele of *RGS1*^*cer7*^ carrying the identified SNP, including 600 bp downstream sequence (3’-UTR), was amplified using allele-specific primers (P1 and P2). The 5’ end of primer P1 is 97 bp upstream of the SNP. The 5’ end of primer P2 included an extension overhanging sequence complementary to the M13F primer site. A 1.4 kb of DNA fragment was amplified from the right flank of the *RGS1* gene from *cer7* using allele-specific primers (P3 and P4). An overhanging sequence was added to the 5’ end of primer P3, complementary to the M13R sequence. The 5’ end of the right flank is 100 bp downstream of the 3’-UTR. Isolated genomic DNA of the *cer7* strain was used as template for the PCR. After amplification the two initial PCR products were mixed with *HY* and *YG*, respectively, and used as templates for a second round of PCR. During the second round PCR, primers P1 and *HY* split were used to amplify a 4.2 kb length fragment, and primers P4 and *YG* split were used to amplify a 2.1 kb length fragment. These products were gel-purified, mixed in equimolar concentrations and transformed into the *Mep2-GFP* strain of *M.oryzae*. The fragments were used to replace the original wild type *RGS1* allele, based on three crossover events (7). The joined *HYG* allele confers hygromycin B resistance and allowed transformants to grow on CM plates supplemented with hygromycin B. We obtained 25 hygromycin B resistant transformants, isolated genomic DNA from these and performed PCR to amplify the fragment from 5’ of left flank of *RGS1* gene locus to 3’ of right flank, using primers P1 and P4. Targeted allelic replacement transformants had an additional 1.5 kb sequence, generating an 8.0kb amplicon, while unsuccessful transformants still contained the wild type *RGS1* allele on a 6.5 kb amplicon. Transformants were selected on complete medium (CM) (1) supplemented with hygromycin B (200 μg mL^−1^) and assessed by PCR to amplify the specific allele which was then sequenced to ensure the *cer7* SNP was present. All mutants and transformants generated in this study are listed in Table S2.

### UV mutagenesis of *M. oryzae*

Conidia of *M. oryzae* were harvested from a CM plate culture using a sterile plastic spreader with 2 mL sterile water. Conidial suspensions were generated in 200 μL sterile distilled water containing 0.1% Tween 20, at a concentration of 5 ×10^5^ conidia mL^−1^ before spreading onto the surface of modified CM agar containing 1% sorbose. The Petri dish was exposed without its lid under UV light at 254 nm using a UV crosslinker (Stratagene) in the dark. A calibration test was first performed to estimate the time necessary to result in either a 40% or 90% kill rate. After UV mutagenesis, plates were immediately wrapped in aluminium foil to avoid DNA photo-repair after ultraviolet light-induced DNA damage. Plates were then incubated at 26°. After 2-3 days, the foil was removed and the plates were grown for a further 4-5 days. Putative mutants were selected based on expression of Mep2-GFP using an epifluorescence stereo microscope Leica M205 FCA. The candidate mutants were then collected using a sterile needle onto fresh CM agar and incubated for 5 days. To perform single spore isolation for each potential candidate mutant, a conidial suspension from each plate was sprayed onto 4% water agar and four single conidia were individually removed using a sterile mounted needle. The resulting single conidia were transferred onto 48 well plates containing CM. The purified mutant strains were then screened for expression of Mep2-GFP as described above and stored as filter paper stocks at −20°C.

### Bulked segregant analysis, SNP calling and alternative splicing analysis

Bulked segregant analysis (BSA) was performed, as previously described with minor modifications (58). Briefly, the *cer7* mutant strain (*Mat1-1*) and wild-type strain TH3 (*Mat1-2*) were grown at 24°C for 7 days on oatmeal agar medium, and then at 18°C until flask-shaped perithecia were visible. After 4-6 weeks growth, mature perithecia were removed and transferred onto 4% water agar medium. The collected perithecia were crushed with a sterile needle to release asci. Mature asci were removed and ascospores dissected under a Leica M205 FCA stereo microscope using a micro-manipulator (Singer Instruments). Collected ascospores were transferred individually to a 48-well plate containing CM and incubated for 5 days (59). Progeny were next screened by phenotypic assessment of fluorescence. Genomic DNA was then extracted from ascospore progeny, as described previously (54). Equal amounts of DNA from each progeny (100ng per progeny) were then bulked. DNA samples were bulked into two samples: wild-type progeny showing low expression of Mep2-GFP, and mutant progeny showing highly expressed Mep2-GFP. The two samples were sequenced using an Illumina HiSeq 2500 generating 125-bp paired-end sequences (University of Exeter Sequencing Service). Low-quality reads were filtered using Trimmomatic (60). SNP calling analysis was performed, as described previously with minor modifications (58). Filtered reads were mapped to the *M. oryzae* 70-15 reference genome (fungi.ensembl.org/Magnaporthe_oryzae/) using Bowtie2 with default parameters (61). Varscan (62) was then used to discover variants by inputting BAM files generated from alignment of reads of the genome sequence of the *cer7* mutant, pooled BSA samples, and the parental Mep2-GFP strain of Guy11 used for mutagenesis. To discover SNPs, filtering was applied based on a minimum read depth of 50 and minimum base identity of 95%. The Integrative Genome Viewer (IGV) (63) was used to manually inspect variants from read alignment for evidence of mutations by input of corresponding bam files.

For alternative splicing analysis, BAM files were generated using RSubread (v.2.10.5) with default settings (64). Putative alternative splicing events of *RGS1* were analyzed based on transcript read densities and visualized using IGV software (63). Peptide sequences of all coding genes in version 8.0 of *M. oryzae* genome sequence were used to predict effectors by EffectorP-2.0 as described (25). Verification of the insertion site of *MEP2-GFP* at a single site was verified by whole genome assembly. Raw reads were aligned to the full plasmid sequence of pCB1532 using Bowtie2 v2.2.9. The sorted BAM file was loaded into the IGV genome viewer to visualize the coverage of raw reads across the Mep2-GFP sequence. The resulting reads aligned to the plasmid and flanking region of the plasmid insertion site were extracted. The total raw reads were realigned against the extracted reads using Exonerate v2.2.0 (32), showing at least 50 bp alignment without any gap, representing reads located at the junction between the inserted plasmid and *M. oryzae* genome. The resulting reads were then aligned against *M. oryzae* genome using Exonerate v2.2.0 and the location of insertion of plasmid visualized via IGV to confirm that the reads aligned to a single locus in Chr1 as a single copy insertion.

### RNA extraction, RNA-seq analysis and quantitative real-time PCR

Conidia from *cer7*, *Δrgs1*, and Guy11 were harvested from 7-day old CM agar plates. Total RNA was extracted from conidia using the Qiagen RNeasy Plant Mini kit according to the manufacturer’s instructions. RNA-seq libraries were prepared using 10 μg of total RNA for Illumina HiSeq 2500 generating 150-bp paired-end sequences (Novogene, Beijing). Low quality reads were filtered by Trimmomatic v0.32 prior to the alignment against version 8.0 of *M. oryzae* genome sequence using bowtie2 (version 2.3.4.3). Quality of alignment was assessed by qualimap2 (version 2.2.1). Pearson correlation and principal component analysis was performed to determine correlation between the samples. The sva package was used to remove batch effects (65). Differentially expressed genes were determined by DESeq2 through a threshold of log2FC and adjusted p-values (p<0.05). for enrichment analysis, Clusterprofiler was used to performed KEGG metabolic pathway enrichment analysis (66). For qRT-PCR, an Affinity Script QPCR cDNA Synthesis Kit was used to generate cDNA, according to manufacturer’s instructions. Briefly, a reaction premixed with 10 μL First-Strand Master Mix, 3μL of oligo(dT) primer, 1 μL of AffinityScript RNase Block enzyme mixture, and 3 μg total RNA, was prepared and incubated at 25 °C for 5 min. The sample was incubated at 42 °C for 15 min to allow cDNA synthesis, before further incubation at 95 °C for 5 min to terminate the cDNA synthesis reaction. Quantitative real-time PCR (qPCR) analysis was conducted using the Stratagene Mx3000TM Real-Time PCR. Each reaction contained 1.25 μL of 606 a 1:5 (v/v) dilution of cDNA, 0.2 μM of each primer and 1X SYBR® Premix Ex Taq™ (Tli RNase H plus, RR420A, Takara) in a total reaction volume of 12.5 μL. PCR conditions were: 1 cycle of 1 min at 95°C; 40 cycles of 5 s at 95°C and 20 s at 60°C; and a final cycle of 1 min at 95°C, 30 s at 58°C and 30 s at 95°C for the dissociation curve. For measurement of the fold change, an efficiency corrected calculation model was made using the formula ((target) ΔCt target (control – sample)) / (housekeeping) ΔCt housekeeping (control – sample) (67).

### Western blot analysis

To test *RGS1* levels in vegetative and invasive growth, total protein was extracted from mycelium and infected rice leaf sheath using *M. oryzae* strains *ToxAp:GFP* (Control) and *RGS1p:GFP-RGS1*. For vegetative growth, mycelium was prepared from CM shake cultures (125 rpm) at 24°C for 48 h. Mycelium was then filtered, washed in distilled water and frozen in liquid nitrogen. For invasive growth, leaf sheath inoculation assays were performed, as previously reported (68) and frozen in liquid nitrogen after 32 hpi. Frozen samples were ground to fine powder in liquid nitrogen using a pestle and mortar. The powder was mixed with 2 times weight/volume ice-cold extraction buffer (10% glycerol, 25 mM Tris pH 7.5, 1 mM EDTA, 150 mM NaCl, 2% w/v PVPP, 10 mM DTT, 1× protease inhibitor cocktail (Sigma), 0.1% Tween 20 (Sigma)), centrifuged at 10,000g at 4°C for 10–20 min, and the supernatant passed through a 0.45 μm Minisart® syringe filter. Proteins were separated by SDS-PAGE and transferred onto a polyvinylidene difluoride (PVDF) membrane using a Trans-Blot turbo transfer system (Bio-Rad). PVDF membrane was blocked with 2% bovine serum albumin (BSA) in Tris-buffered saline and 1% Tween 20 (Sigma). GFP detection was carried using a GFP (B2):sc-9996 horseradish peroxidase (HRP)-conjugated antibody (Santa Cruz Biotechnology, Santa Cruz, CA). An anti-actin antibody was used as the loading control for fungal and plant protein. Pierce ECL Western Blotting Substrate (Thermo Fisher Scientific) was used for detection. Membranes were imaged using ImageQuant LAS 4000 luminescent imager (GE Life Sciences).

### Yeast Transactivation Assay and Yeast One-Hybrid Analysis

The transcriptional activity and DNA-binding ability of N-Rgs1 and Rgs1 were tested in *Saccharomyces cerevisiae* strain Y2HGold. Sequences encoding N-Rgs1 and C-Rgs1 were amplified from cDNA derived from total RNA of Guy11 conidia. Fragments were cloned into pGBKT7 (BD) and pGADT7 (AD) vectors, respectively, by in-fusion cloning (In-Fusion Cloning kit; Clontech Laboratories). pGBKT7(BD)-N-Rgs1, pGBKT7(BD)-Rgs1 were used to test transactivation activity, and pGADT7(AD)-N-Rgs1, pGADT7(AD)-Rgs1 were used to test DNA-binding ability. The empty vectors were co-transformed and used to test the transcriptional activation as a negative control. The vectors pGBKT7-53 and pGADT7-T were used as the positive control. Empty un-linearized bait vector pGBKT7 and prey vector pGADT7 were used as negative controls. BKT variants and GAD variants were co-transformed into chemically competent Y2HGold yeast cell following the manufacturer’s manual. Successfully transformed yeast colonies were inoculated in 2mL of liquid SD/-Leu/-Trp selection media for overnight growth at 30°C. The yeast samples were then used to produce dilutions of OD_600_ 1, 1^−1^, 1^−2^, 1^−3^, and 1^−4^, respectively. Cell suspensions in 10μL droplets of each dilution were spotted on a SD/-Leu/-Trp/-His to determine the transactivation activity and DNA-binding ability, after 72 h growth at 30°C.

For yeast one-hybrid experiments, the Matchmaker Gold Yeast One-hybrid System kit (Takara Bio, USA) was used according to manufacturer’s instructions. Briefly, a 1 kb fragment upstream of the *MEP2* gene was prepared as the bait sequence and ligated into digested vector pAbAi, which carries the bAr gene (AUR-1C) that confers resistance to AbA (Aureobasidin A) (Takara Bio, USA). The p-AbAi-*MEP2* vector, and positive control vector p53-AbAi were digested with BbsI, and then transformed into yeast strain Y1HGold (69), respectively. Y1HGold [*MEP2p* / AbAi] and Y1HGold [p53 / AbAi] was screened by growing on SD/-Ura. The minimum inhibitory concentration of AbA (200 ng/mL) for the bait strain was determined by growing on SD/-Ura/AbA medium. The cDNA sequence encoding N-Rgs1 was amplified from total RNA by RT-PCR using primers with overhanging sequences the prey vector pGADT7. The plasmid pGADT7-was then transformed into bait yeast strain Y1HGold [*MEP2p*/AbAi], and Y1HGold[p53/AbAi], respectively. The yeast samples were used to produce dilutions of OD600 1, 1^−1^, 1^−2^, 1^−3^, and 1^−4^, respectively. SD/-Leu/AbA^200^ medium was used to test the *MEP2p*/N-Rgs1 interaction.

## Acknowledgments

This project was supported by a European Research Council Advanced Investigator award (to NJT) under the European Union’s Seventh Framework Programme FP7/2007-2013/ERC Grant Agreement 294702 GENBLAST, a grant to NJT from The Gatsby Charitable Foundation, and a Halpin Scholarship to BT.

## Author Contributions

Author contributions: B.T. and N.J.T. designed research; B.T., X.Y., A.J.F., C.M., and N.C.M. performed experiments. B.T., and D.M.S. performed analysis of data. B.T., L.S.R., and N.J.T. wrote the paper.

The authors declare no conflict of interest.

Data deposition: RNA-seq data of *M. oryzae* and the genome sequence of *cer7* mutant strain reported in this paper were deposited in the European Nucleotide Archive - EMBL-EBI database under accession number: PRJEB45710.

---

**Fig. S1.**
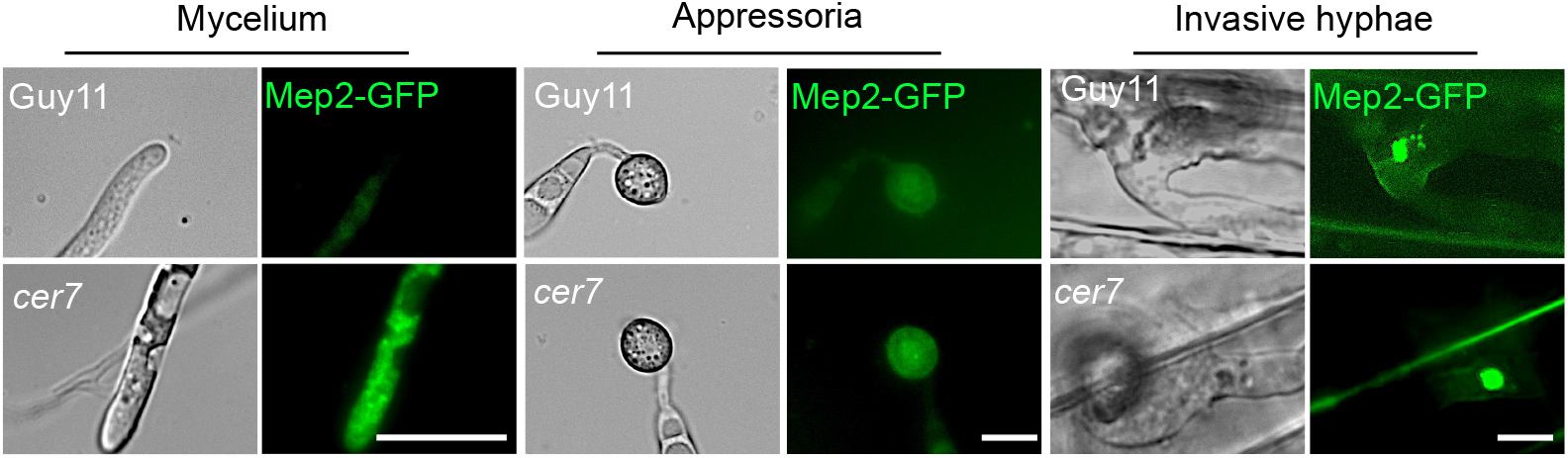
Expression of Mep2-GFP in the *cer7* mutant strain is constitutively activated during different development stages including mycelium, appressoria, and invasive hyphae. Micrographs showing fluorescence of Mep2-GFP in Guy11 and the *cer7* mutant, in mycelium, appressoria, and invasive hyphae. Mycelium was imaged after 24 hours growing in liquid CM culture. The image for appressorium development was captured 8hpi on a hydrophobic glass coverslip. Invasive hyphae were imaged 32hpi on CO39 rice leaf sheath inoculated with conidial suspensions at 1×10^4^/mL (Scale bar, 10 μm).

**Fig. S2.**
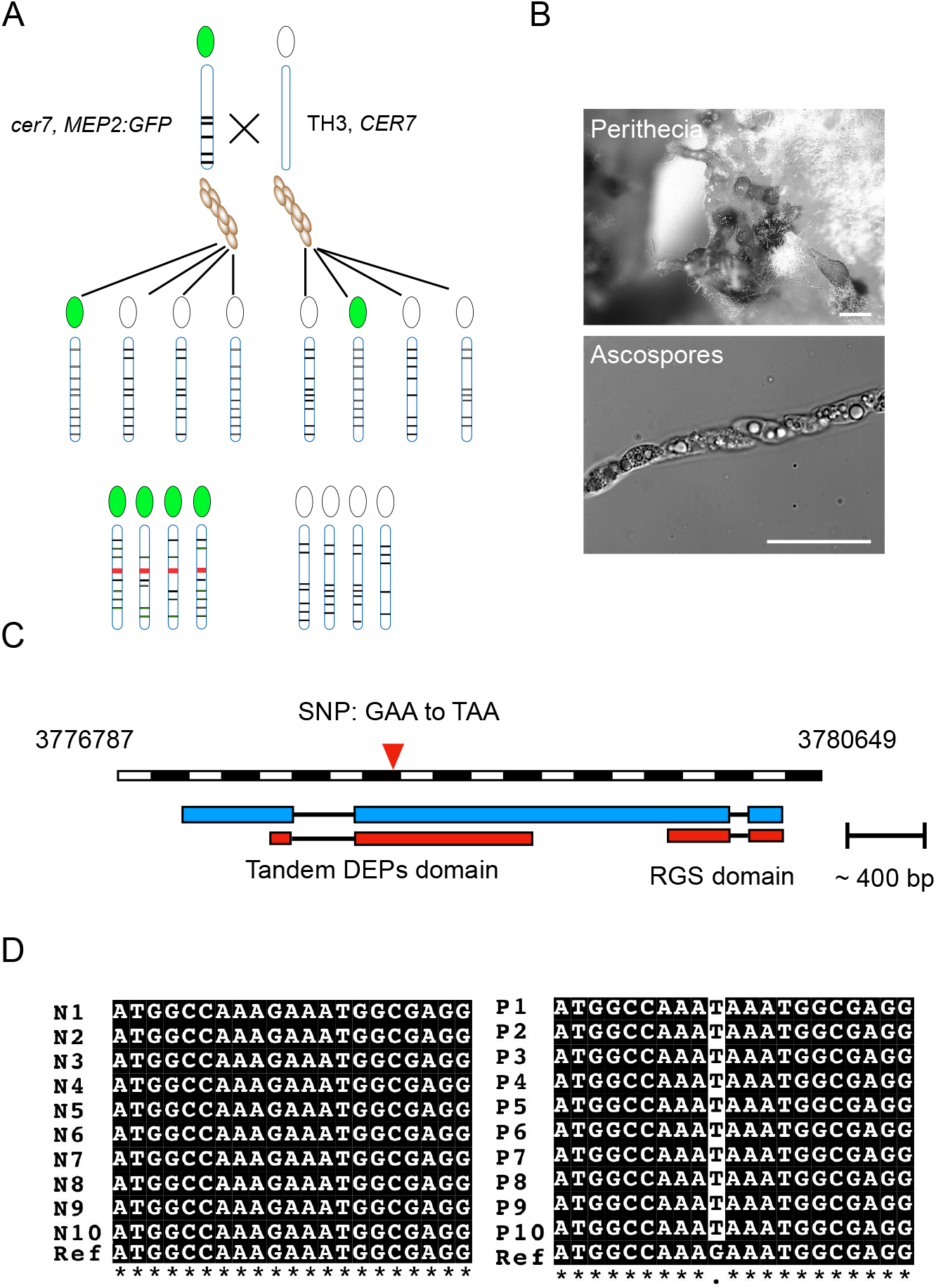
Bulk segregant analysis defined a region of chromosome 2 showing very strong linkage to *cer7*. (***A***) Diagram to show bulked segregation analysis strategy to identify the mutant allele causing constitutive expression of *MEP2* in conidia. The *cer7* mutant (*Mat1-2*) and TH3 (*Mat1-1*) were used in sexual crosses by incubation together on oatmeal agar. After 4-6 weeks, perithecia developed and were collected and manipulated to release asci containing ascospore progeny. (***B***) Micrographs to show the perithecia and eight-spored ascus obtained from crossing *cer7* strain and TH3. (Scale bar, 10 μm). Images were captured after 8 weeks growth on oatmeal agar. (***C***) Overview of the genomic region of the *RGS1* locus to show transcript structures, and location of detected SNP in the highest linkage region identified by BSA. (***D***) Sequence analysis of amplicons of the *RGS1* coding sequences from 20 ascospore progeny. N1 to N10 are amplicons from progeny showing low Mep2-GFP fluorescence (wild-type), and P1 to P10 represent amplicons of progeny showing high expression of Mep2-GFP (*cer7*). “Ref” shows the DNA sequence of *RGS1* in the *M. oryzae* reference genome.

**Fig. S3.**
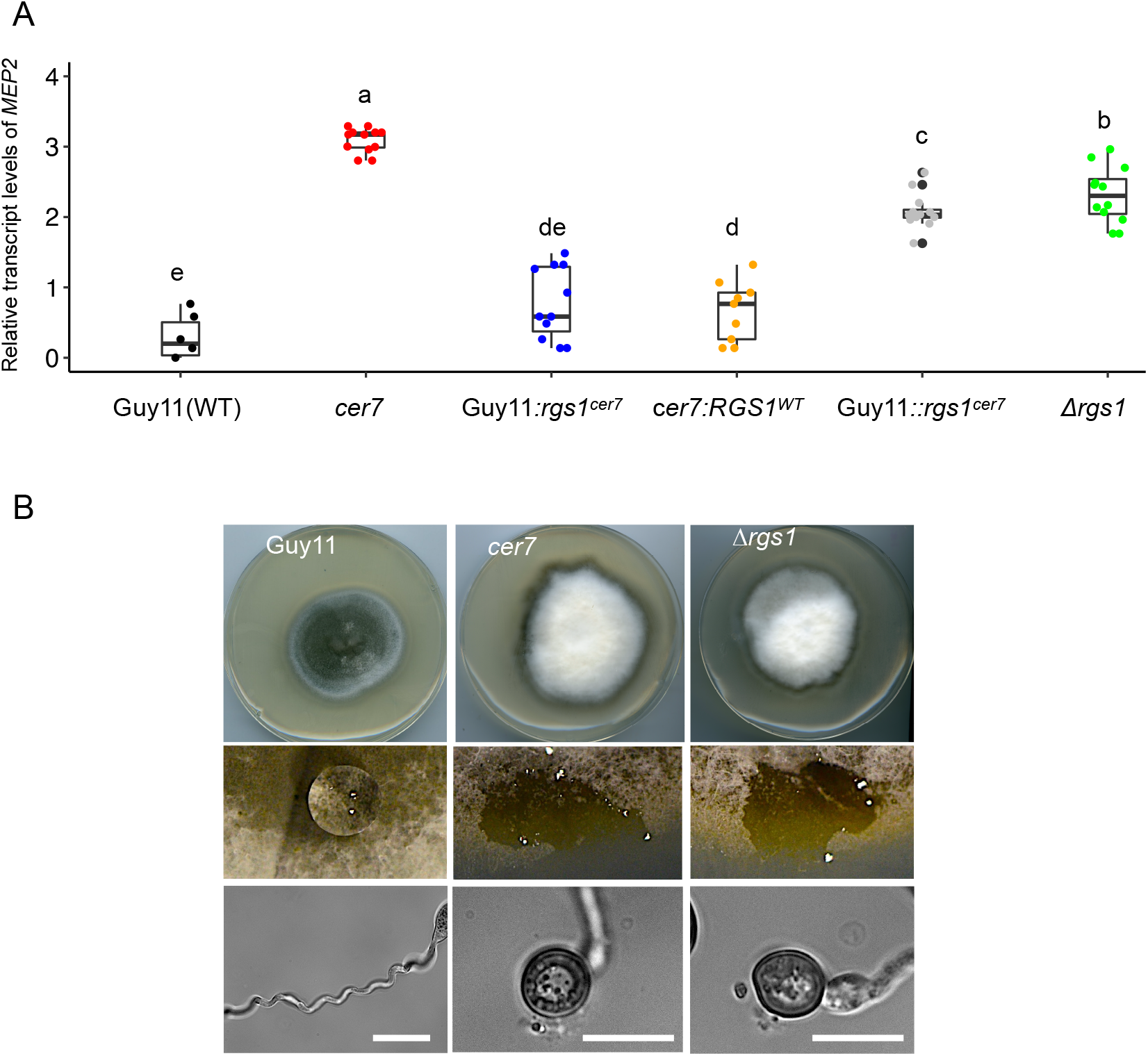
Complementation analysis of the *cer7* mutant with *RGS1*. (***A***) Box plot to show fold change in relative transcripts of *MEP2* from mycelium of the wild type Guy11 (Black), *cer7* mutant (red), Guy11:*rgs1*^*cer7*^ ectopic transformant (blue), *cer7:RGS1*^*WT*^ complemented transformant (orange), Guy11::*rgs1*^*cer7*^ allelic replacement mutant (grey) *Δrgs1* mutant (green). Expression was determined relative to the fungal actin gene. Letters refer to significant differences determined by One-way ANOVA tests (p<0.05, Duncan test). (***B***) Upper panel Images to show the colony morphology of wild type Guy11, compared to *cer7* and Δ*rgs1* mutants. Both *cer7* and Δ*rgs1* form white fluffy colonies when grown on complete medium. Mid panel images show droplets of water placed on the surface of plate cultures of Guy11, *cer7* and Δ*rgs1* mutant strains. Mutants show an easily-wettable phenotype. Photographs were taken 24hpi. Lower panel micrographs show appressorium development by *cer7* and Δ*rgs1* mutants on a non-inductive hydrophilic surface. (Scale bar, 10 μm).

**Figure S4.**
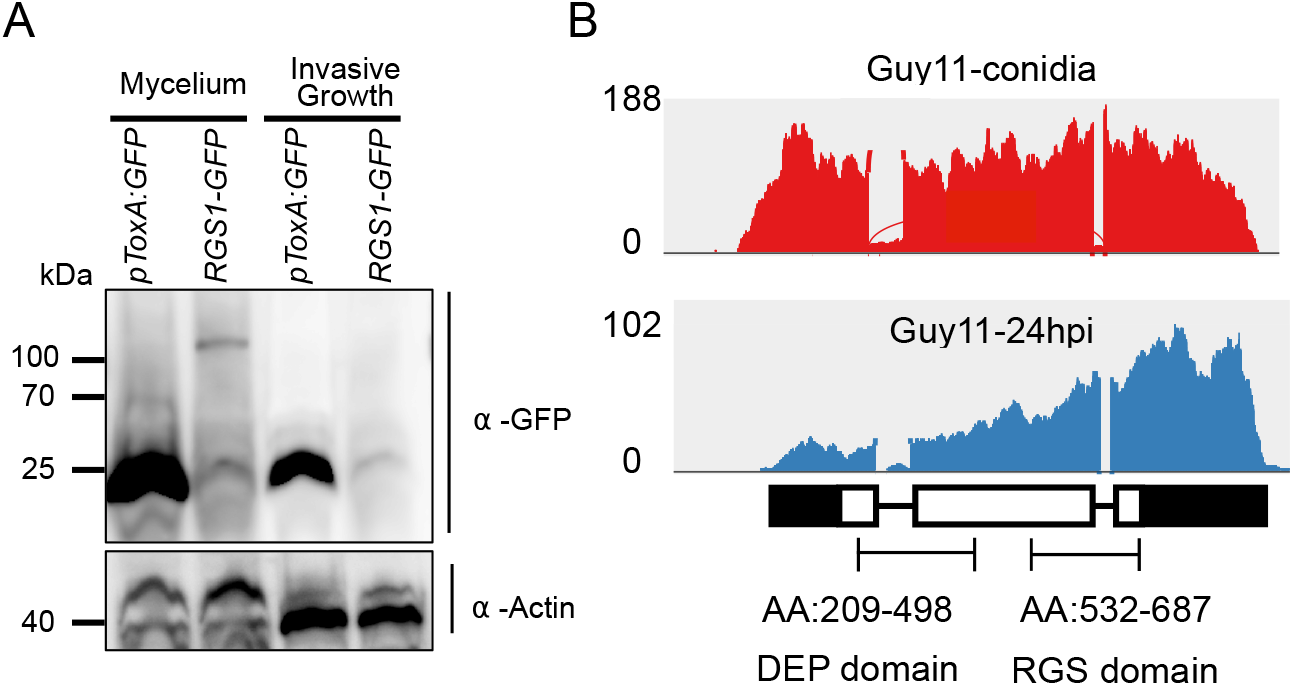
Expression analysis of *RGS1*. (***A***) Western blot showing abundance of Rgs1-GFP detected with anti-GFP antibody in mycelium and invasive hyphae (IH). Total protein extracted from mycelium of Rgs1-GFP after growing in liquid CM culture for 48 hours. Lysates were resolved by SDS-PAGE and analysed by western blot using antibodies against GFP. Detection using anti-actin was used as the loading control for fungal protein. Molecular mass standards in kDa are indicated on the left. (***B***) Sashimi plots showing mRNA transcript reads of *RGS1* in Guy11, as conidia at 0hpi (red), and during infection at 24hpi (blue). The bar plot demonstrated the depth of reads aligned to corresponding regions of the gene. RNA-seq from libraries of total RNA isolated from leaf-drop infection inoculated by wild-type Guy11 were used. The numbers on the left panel indicate the coverage of reads detected after alignment and visualized in IGV.

**Fig. S5.**
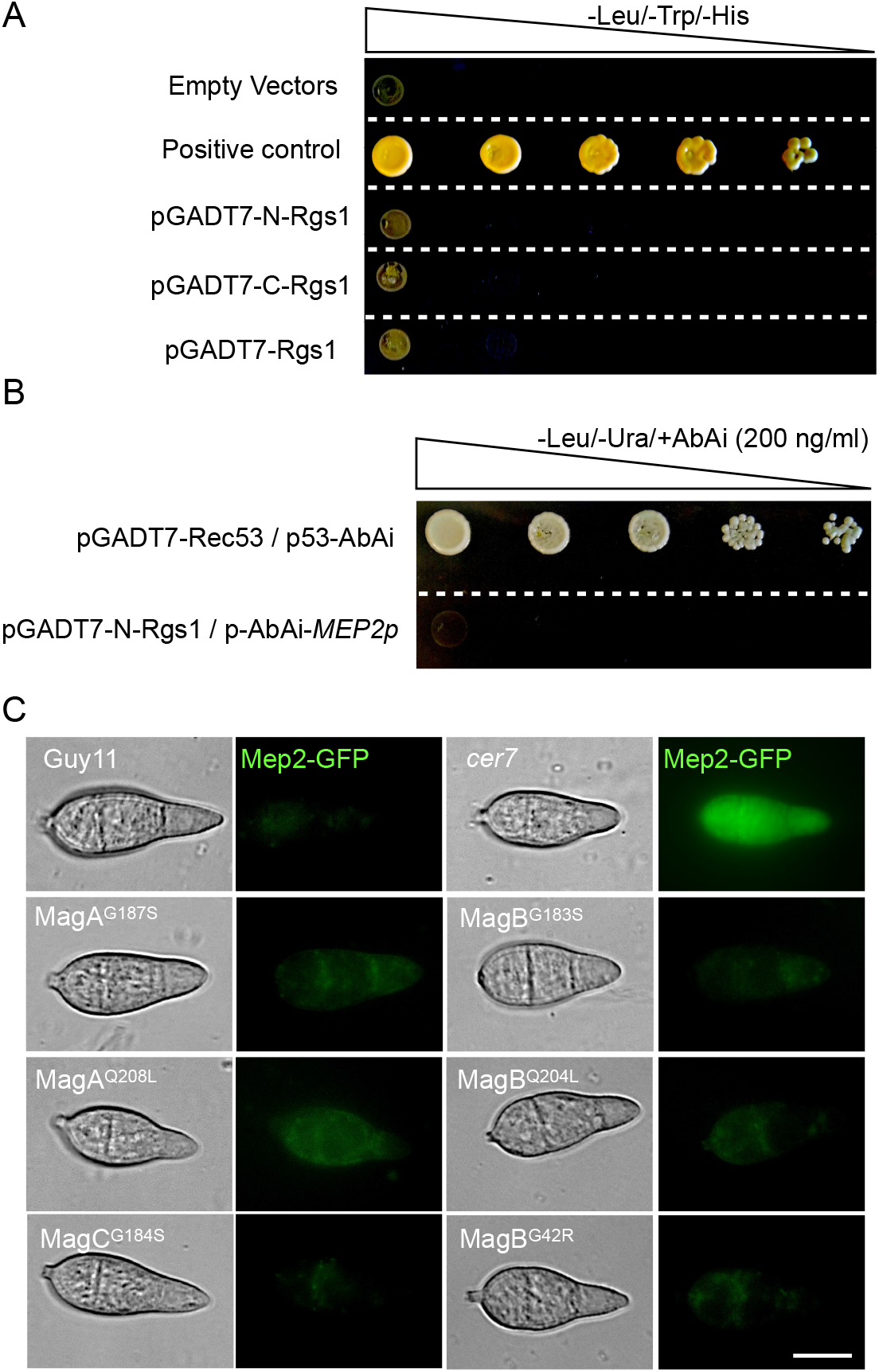
N-Rgs1 does not possess DNA-binding ability or bind to the *MEP2* promoter. (***A***) DNA-binding assay of N-Rgs1, C-Rgs1, and full length Rgs1 in yeast cells. Co-transformation of Y2HGold strains was carried out with bait (BD) and prey (AD) vectors in the following combinations; pGBKT7 / pGADT7-N-Rgs1, pGBKT7 / pGADT7-C-Rgs1, pGBKT7 / pGADT7-Rgs1, and positive control (pGBKT7-53 and pGADT7-T) along with empty vectors. Cells were grown on double drop out quadruple dropout medium. Images are representative of two biological replicates. (***B***) Yeast-one-hybrid assay to determine the interaction between Rgs1 and upstream sequences of the *MEP2* gene. The Y1HGold strain containing p-AbAi-*MEP2p* was transformed by introducing plasmid pGADT7-Rgs1^1-498^ and used to test growth on medium with -Leu/-Ura, supplemented with 200 ng/mL Aureobasidin A (AbA). (***C***) The repression of *MEP2* expression by Rgs1 occurs in a G-protein signalling-independent manner. Micrographs showing expression of Mep2-GFP in conidia of Guy11, *cer7*, MagA^G187S^, MagB^G183S^, MagA^Q208L^, MagB^Q204L^, MagC^G184S^, and MagB^G42R^. MagA^G187S^, MagB^G183S^, and MagC^G184S^. Images are representative at least 10 transformants examined for each experiment. Conidial suspensions from each strain were harvested from colonies after 5 days growth on CM and immediately visualized using epifluorescence microscopy. (Scale bar, 10 μm).

**Fig. S6.**
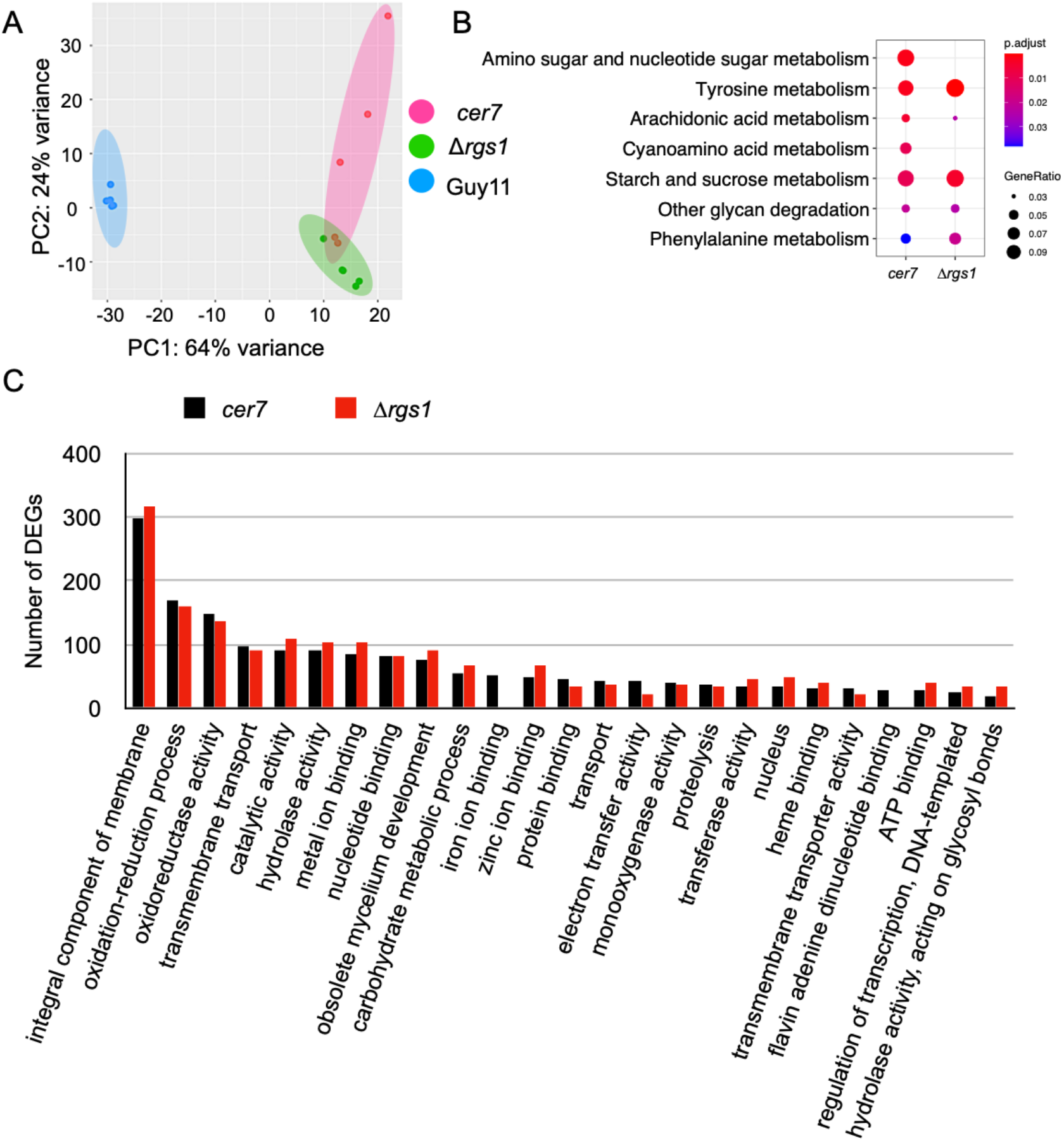
RNA-seq analysis of *cer7*, *Δrgs1* and Guy11. (***A***) Plot showing Principal Component Analysis (PCA) to determine the distance between samples analysed by RNA-seq. Genes with raw reads counts of less than 10 were discarded, and remaining read counts normalized as FPKMs and used to perform analysis to generate Principal Components (PCs). The x-axis shows PC1, and the y-axis show PC2. Red dots represent samples from the *cer7* mutant, blue dots represent Guy11, and green dots represent the Δ*rgs1* mutant. (***B***) Dot plot showing the comparative metabolic pathway enrichment analysis of DEGs identified from RNA-seq of *cer7* and Δ*rgs1*. The size of dots indicates the ratio of the number of DEGs in the pathway / the number of total DEGs. (***C***) Bar chart showing the top 25 biological processes enriched in DEGs in *cer7* and *Δrgs1* mutants compared to Guy11, based on Gene Ontology analysis of RNA-seq data. The X-axis shows corresponding functions according to the Gene Ontology annotation of *M. oryzae*. The Y-axis shows the number of DEGs identified from the RNA-seq dataset. Black bar plots represent the comparison between *cer7* and Guy11. Red bar plots represent comparison between *Δrgs1* and Guy11.

**Fig. S7.**
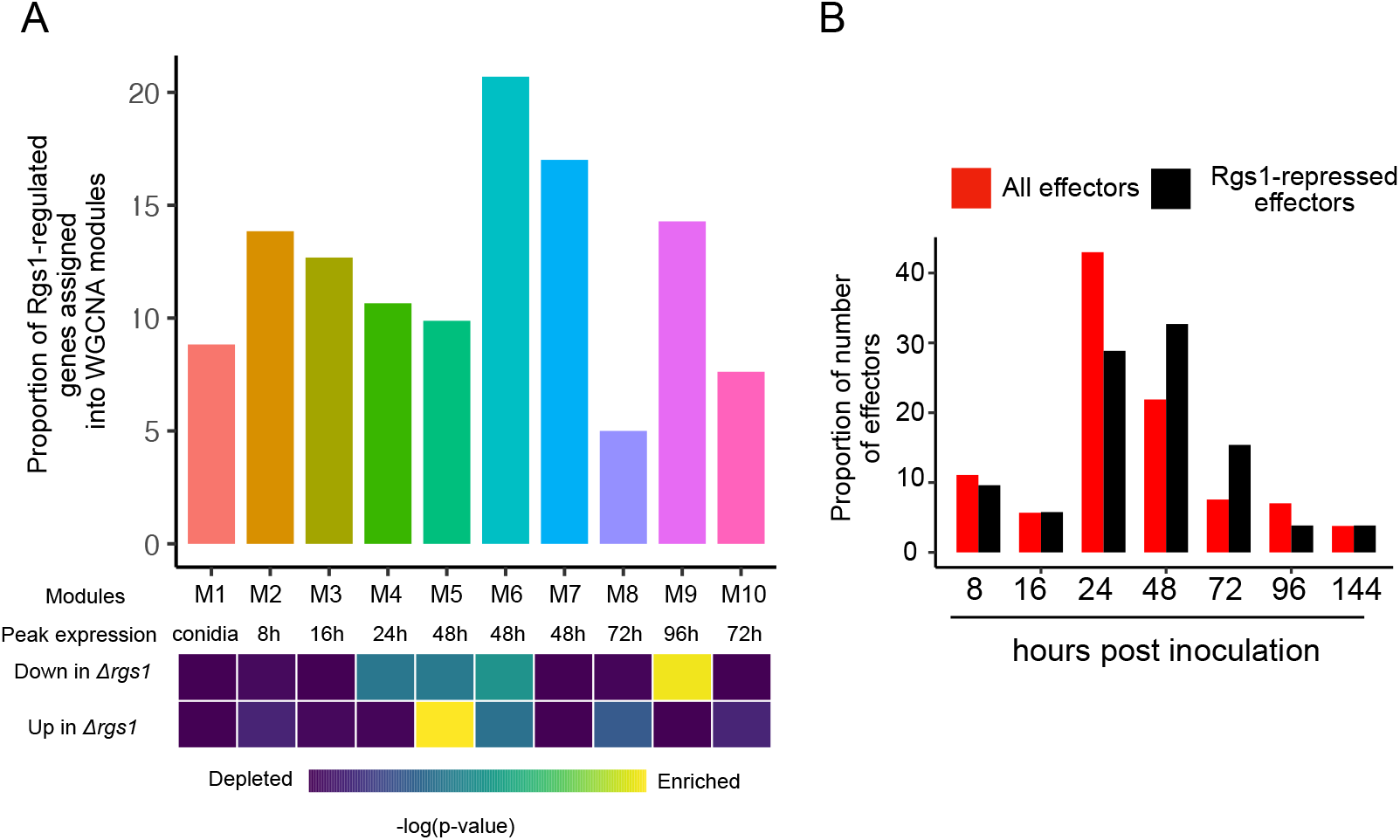
Rgs1-regulated effectors temporally co-expressed during plant infection. (***A***) Bar chart to show the fraction of Rgs1-regulated genes assigned to each WCGNA co-expression module defined in Ref. 1. X-axis shows the time-point of peak expression during plant infection as described previously (25). The heatmap shows enrichment of Rgs1-regulated genes. Enrichment of genes up-regulated in *Δrgs1* conidia was observed in WCGNA modules M4, M5 and M6 which show peak expression at 24-48hpi. Genes down-regulated in *Δrgs1* conidia was observed in M9, which peaks in expression at 96hpi. (***B***). Bar chart to show temporal expression profiles of *M. oryzae* effector-encoding genes (25). Black bars indicate the number of Rgs1-regulated effectors, and red bars the total number of putative effectors expressed at each time point. Rgs1-regulated effectors are over-represented among those peaking in expression at 24h and 48h, consistent with WCGNA co-expression analysis.

**Fig. S8.**
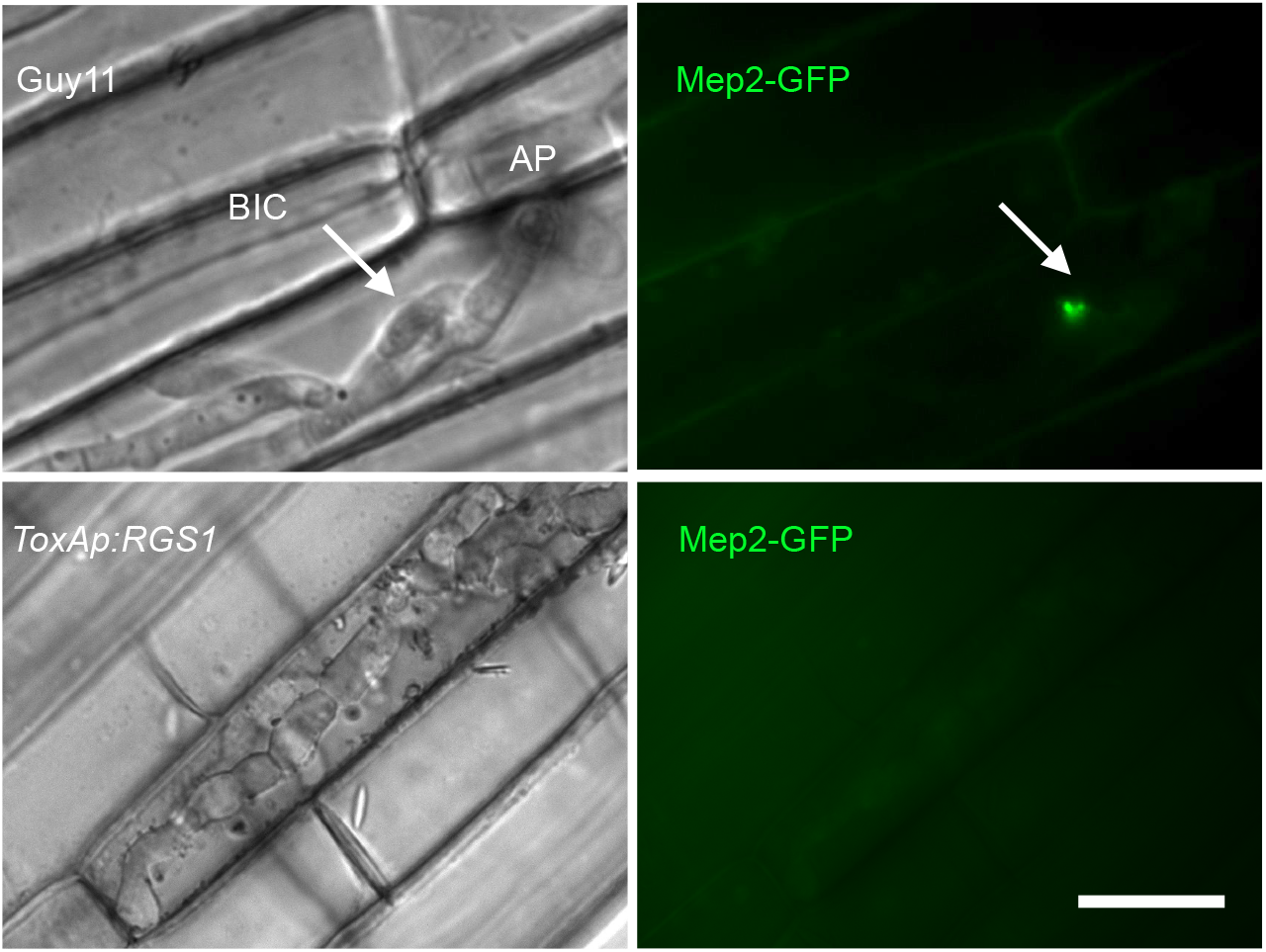
Constitutive Rgs1 over-expression prevents Mep2-GFP expression *in planta*. Micrographs to show expression of Mep2-GFP in the BIC of invasive hyphae of Guy11 and the *ToxAp:RGS1* strain which constitutively expresses *RGS1*. BIC localization was imaged in rice leaf sheath tissue inoculated by conidia of Guy11:Mep2-GFP at 32 hpi. (Scale bar, 10 μm).

**Table S1.**
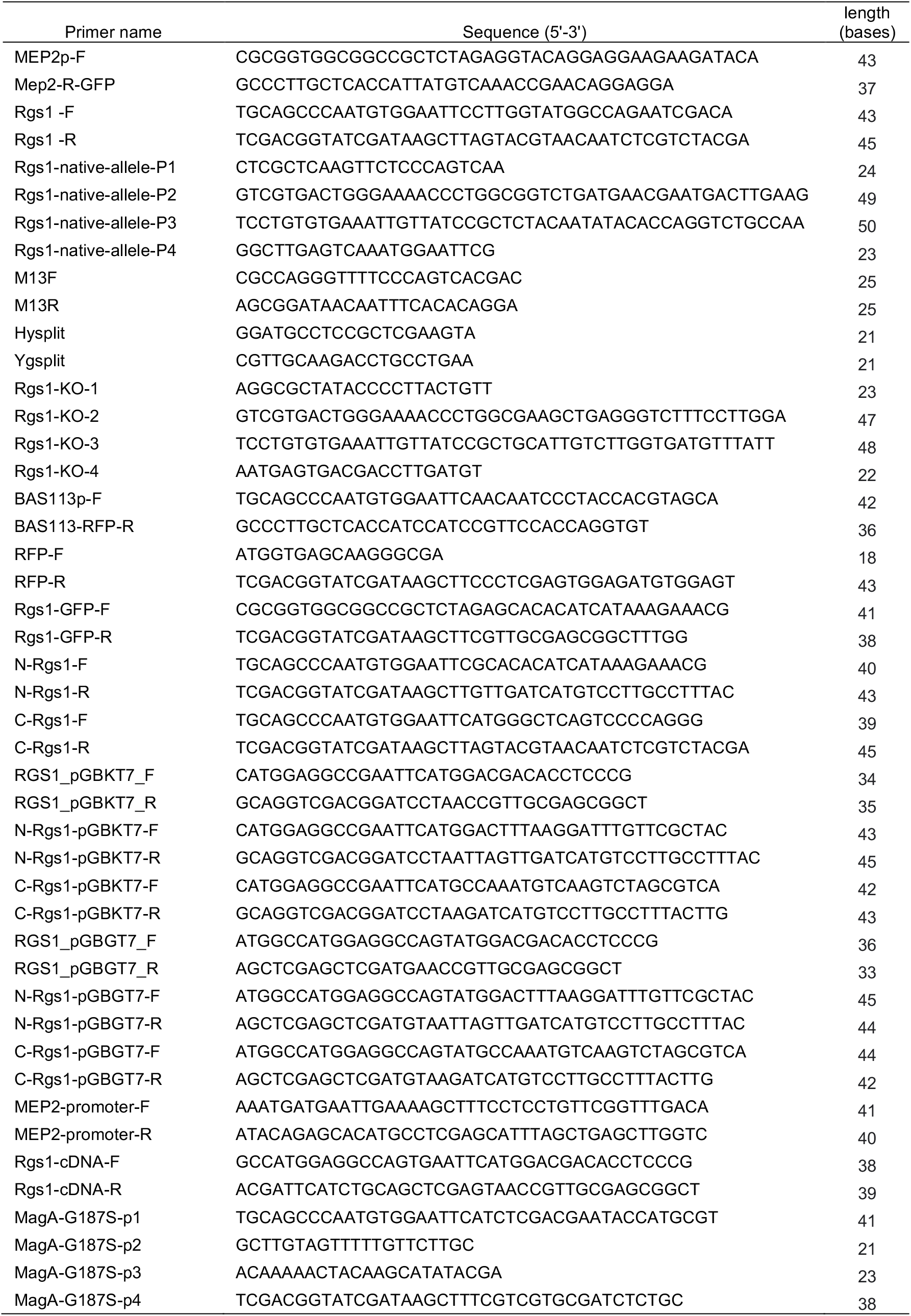
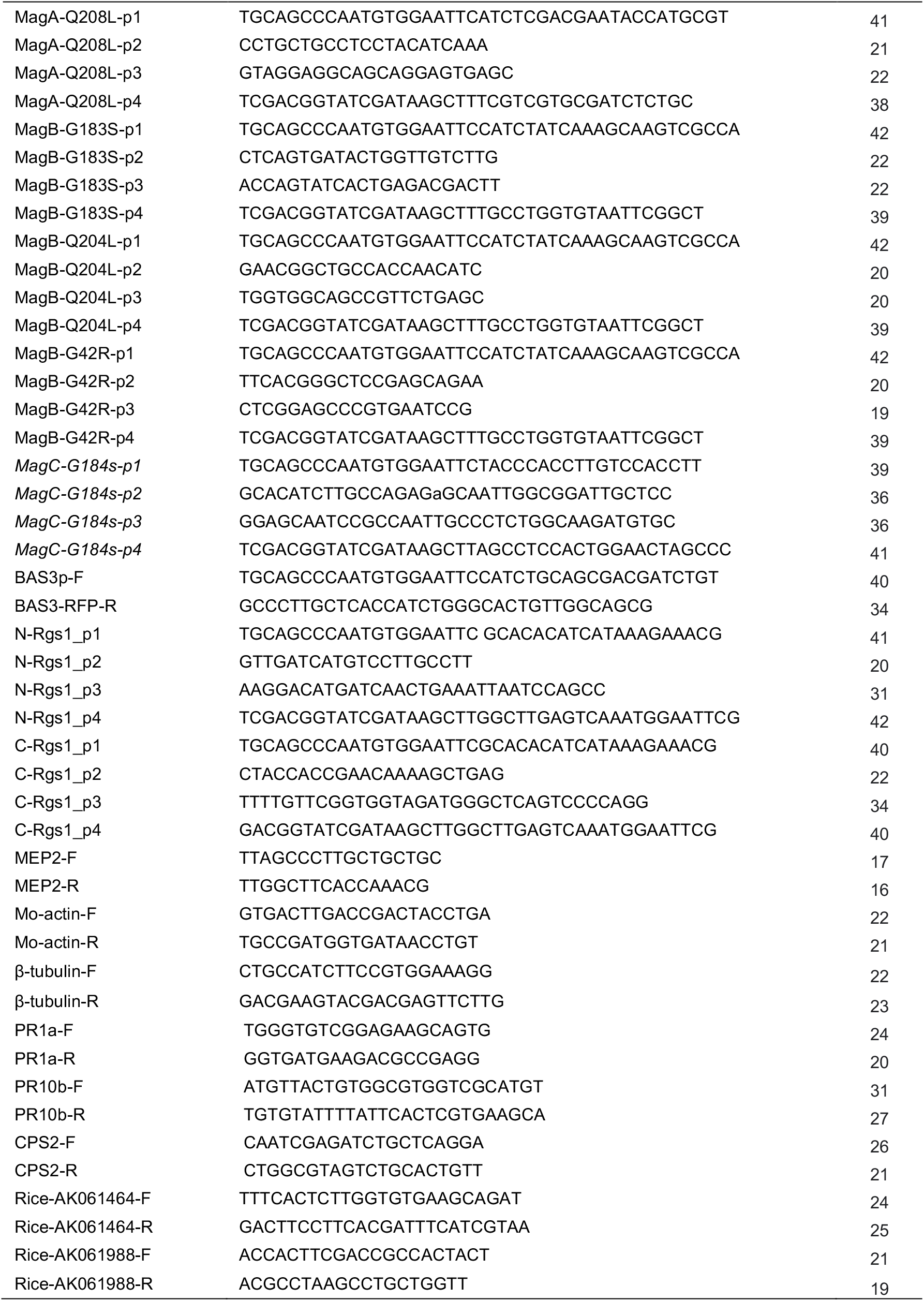
Primers used in this study.

**Table S2.** *Magnaporthe oryzae* strains generated in this study

| Strain Name | Description |
| --- | --- |
| cer7 | Guy11 constitutively expressing Mep2-GFP after UV mutagenesis |
| Mep2-GFP | Guy11 transformed with plasmid pMep2-GFP |
| cer7/Rgs1cer7 | Cer7 transformed with plasmid Rgs1cer7 |
| cer7/Rgs1wt | Cer7 transformed with plasmid Rgs1wt |
| Δrgs1 | targeted gene deletion in Rgs1 locus |
| Rgs1CER7 | Guy11 transformed with plasmid pMep2-GFP and transformed with native allele of Rgs1 carrying the Cer7 mutant |
| Δrgs1/Mep2-GFP | Δrgs1 transformed with plasmid pMep2-GFP |
| Mep2-GFP/rgs1cer7 | Guy11 strain with Mep2-GFP and transformed with plasmid rgs1cer7 |
| pToxA-GFP | Guy11 with constitutively expressed GFP driven by pToxA |
| Rgs1-GFP | Guy11 transformed with plasmid Rgs1-GFP |
| cer7/N-Rgs1 | Cer7 transformed with plasmid N-Rgs1 |
| cer7/C-Rgs1 | Cer7 transformed with plasmid C-Rgs1 |
| MagA-Q208L | Guy11 constitutively expressing Mep2-GFP and transformed with plasmid MagA-Q208L |
| MagA-G187S | Guy11 constitutively expressing Mep2-GFP and transformed with plasmid MagA-G187S |
| MagB-G183S | Guy11 constitutively expressing Mep2-GFP and transformed with plasmid MagB-G183S |
| MagB-Q204L | Guy11 constitutively expressing Mep2-GFP and transformed with plasmid MagB-Q204L |
| MagB-G42R | Guy11 constitutively expressing Mep2-GFP and transformed with plasmid MagA-Q208L |
| MagC-G184S | Guy11 constitutively expressing Mep2-GFP and transformed with plasmid MagC-G184S |
| Cer7/bas113-RFP | Cer7 transformed with plasmid Bas113-RFP |
| Cer7/Bas3-RFP | Cer7 transformed with plasmid Bas3-RFP |
| pToxA-Rgs1-GFP | Guy11 transformed with plasmid pToxA-Rgs1-GFP |

**Table S3.**
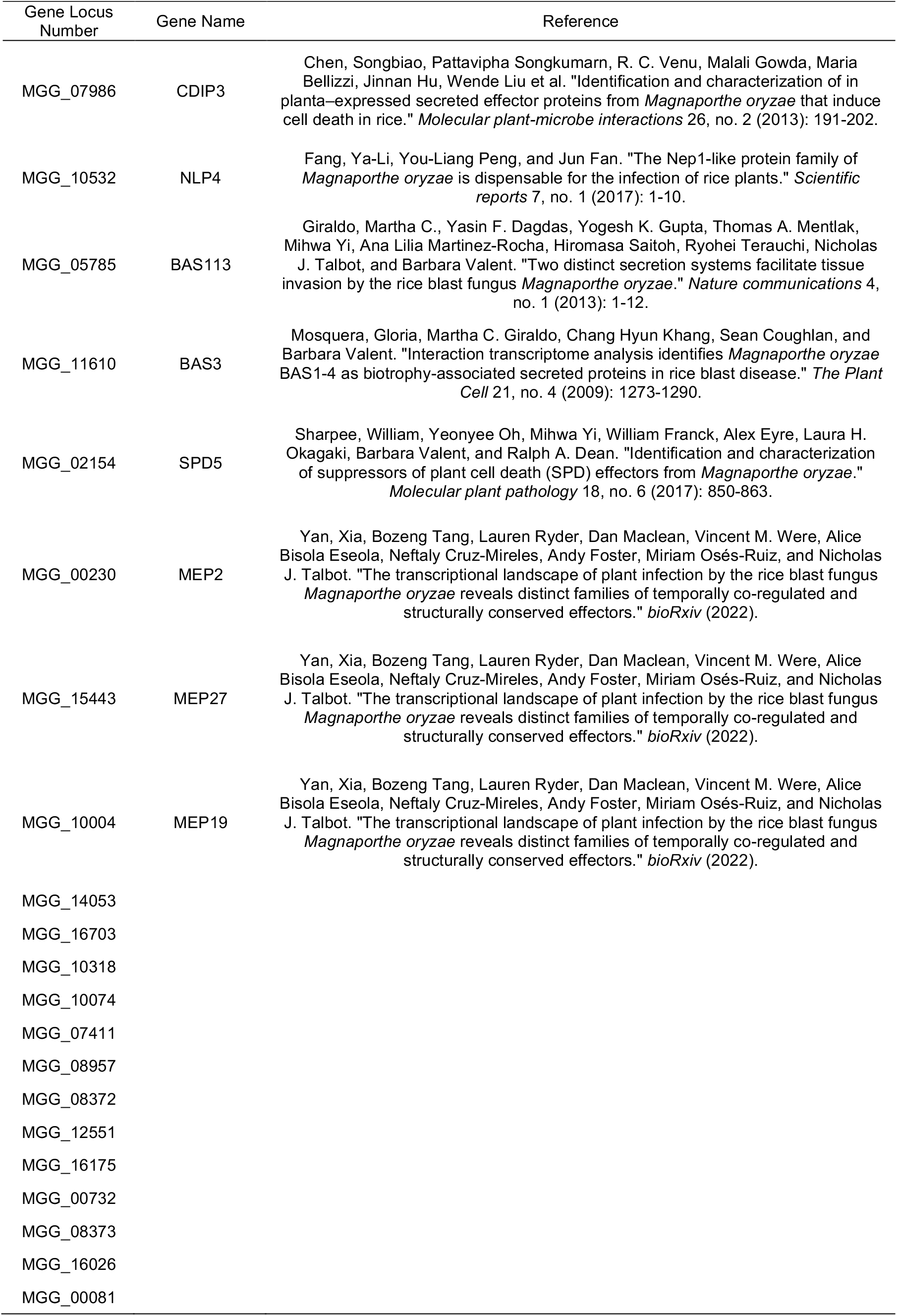
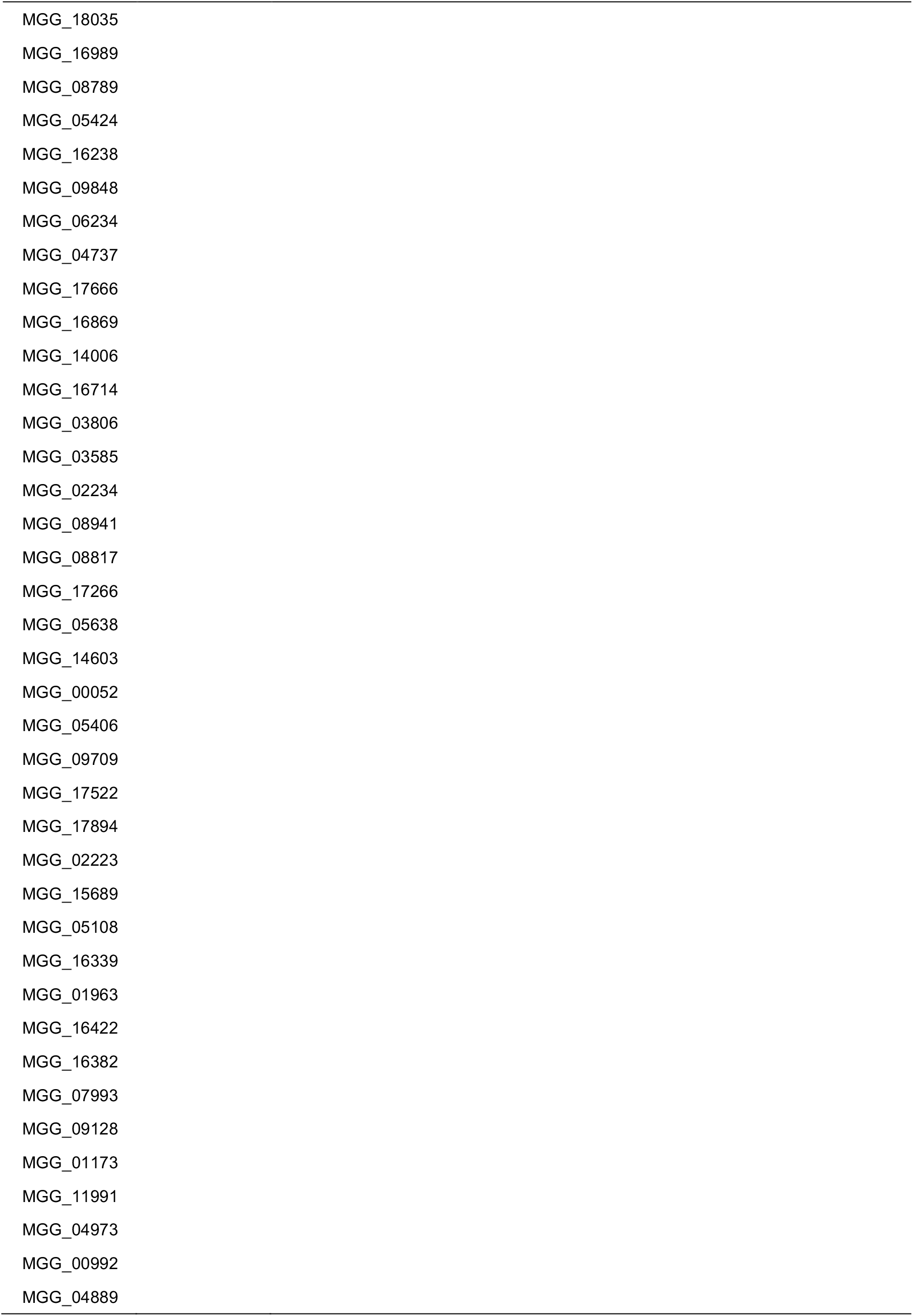
Rgs1-regulated effectors identified in this study

